# Electrostatic encoding of genome organization principles within single native nucleosomes

**DOI:** 10.1101/2023.12.08.570828

**Authors:** Sangwoo Park, Advait Athreya, Gustavo Ezequiel Carrizo, Nils A. Benning, Michelle M. Mitchener, Natarajan V. Bhanu, Benjamin A. Garcia, Bin Zhang, Tom W. Muir, Erika L. Pearce, Taekjip Ha

## Abstract

The eukaryotic genome, first packed into nucleosomes of about 150 bp around the histone core, is organized into euchromatin and heterochromatin, corresponding to the A and B compartments, respectively. Here, we asked if individual nucleosomes in vivo know where to go. That is, do mono-nucleosomes by themselves contain A/B compartment information, associated with transcription activity, in their biophysical properties? We purified native mono-nucleosomes to high monodispersity and used physiological concentrations of biological polyamines to determine their condensability. The chromosomal regions known to partition into A compartments have low condensability and vice versa. *In silico* chromatin polymer simulations using condensability as the only input showed that biophysical information needed to form compartments is all contained in single native nucleosomes and no other factors are needed. Condensability is also strongly anticorrelated with gene expression, and especially so near the promoter region and in a cell type dependent manner. Therefore, individual nucleosomes in the promoter know whether the gene is on or off, and that information is contained in their biophysical properties. Comparison with genetic and epigenetic features suggest that nucleosome condensability is a very meaningful axis onto which to project the high dimensional cellular chromatin state. Analysis of condensability using various condensing agents including those that are protein-based suggests that genome organization principle encoded into individual nucleosomes is electrostatic in nature. Polyamine depletion in mouse T cells, by either knocking out ornithine decarboxylase (ODC) or inhibiting ODC, results in hyperpolarized condensability, suggesting that when cells cannot rely on polyamines to translate biophysical properties of nucleosomes to control gene expression and 3D genome organization, they accentuate condensability contrast, which may explain dysfunction known to occur with polyamine deficiency.

## INTRODUCTION

The nuclear genome is largely partitioned into two domains: the gene-rich and relatively open euchromatin and the gene-poor and relatively compact heterochromatin. With the advent of new technologies such as Hi-C and chromatin tracing, the complex hierarchal organization of the genome is now being appreciated ^1–4^. Each chromosome occupies its own territory in the nucleus, and single chromosomes are partitioned into A and B compartments on a multi-mega base pair scale ^5^. They are further compartmentalized into topologically associated domains (TADs) and loops in a 1 mega to 10 kilo base pair scale ^6,7^. Phase-separation of biomolecules has been suggested as the biophysical basis of many membrane-less compartments in cells ^8–10^, including those of chromatin. Electrostatic interactions, hydrophobic interactions, and cation-pi interactions seem to be important drivers of this phenomenon ^10,11^. Heterochromatin organization has also been explained in terms of chromatin condensation, with either liquid-like ^12–14^ or gel-like ^15^ properties. The heterochromatin is AT-rich and possesses many non-coding repeat sequences. In contrast, highly transcribing genes usually tend to have low AT content ^16–18^. GC-rich CpG islands are frequently found in active promoters where they are largely unmethylated. Conversely, CpG islands on silent promoters tend to be densely methylated ^19,20^. Histone post-translational modifications (PTMs) and histone variants also reflect the functional state of the chromatin ^21,22^.

Although the biological functions of genetic–epigenetic features have been mainly interpreted in the context of interacting partners, such as readers and writers of specific DNA sequences or epigenetic codes ^23–25^, their intrinsic physical properties can also have direct biological implications. DNA sequences, with AT content or even a long poly(dA:dT) tract, have peculiar groove structures and curvatures, which can play special roles in ionic interactions ^26–30^. Histone PTMs could be important modulators for determining the intrinsic properties of nucleosomes ^31^. DNA methylation also modulates ionic interactions between DNAs through changing the groove width of cytosine and guanine pairs ^30,32^. Despite extensive knowledge of genome organization, understanding of the biophysical driving force behind genomic compartmentation remains elusive. In this study, we ask whether the nucleosomes know where to go. That is, do individual nucleosomes in the cell intrinsically encode information important for their participation in large scale organizations such as A and B compartments and in local organizations such as promoters, enhancers and gene bodies? To address these questions, we developed an assay to measure their intrinsic condensability mediated by physiological condensing agents.

## RESULT

### Condense-seq measures native mono-nucleosome’s condensability genome-wide

We used various DNA and nucleosome condensing agents ^33^, including polyamines ^34^, cobalt- hexamine ^35^, polyethylene glycol (PEG) ^36^, calcium ^37^, HP1α and HP1β ^12,38^ to induce condensation of native nucleosomes in vitro. Native mono-nucleosomes were prepared with hydroxy apatite purification ^39^ after in-nuclei micrococcal nuclease digestion of the chromatin, followed by further size-selection to obtain monodisperse samples (Fig. 1a and Extended Data Fig. 1). The monodispersity afforded by mono-nucleosome purification was critical for quantitative comparison among native nucleosomes. The nucleosome condensation experiment was first performed under various concentrations of spermine as a condensing agent (Fig. 1b). Compared to DNA condensation, obtained after deproteinization, nucleosome condensation showed a wider distribution of polyamine concentration required for condensation, which we attribute to histone modifications and variants. Through sequencing the nucleosomes remaining in the supernatant and comparing them with the input control, each nucleosome could be localized along the genome and its survival probability after condensation could be estimated (Extended Data Fig. 2a,b). We defined the “condensability”, the propensity of being incorporated into precipitation, as the negative natural log of the survival probability (Fig. 1a). From this “condense-seq” assay, we can determine genome-wide condensability at the single-nucleosome resolution (Extended Data Fig. 2). First, we focused on nucleosome condensation induced by spermine which is a small biological metabolite and a prevalent polyamine in eukaryote nuclei ^40,41^.

**Fig. 1.**
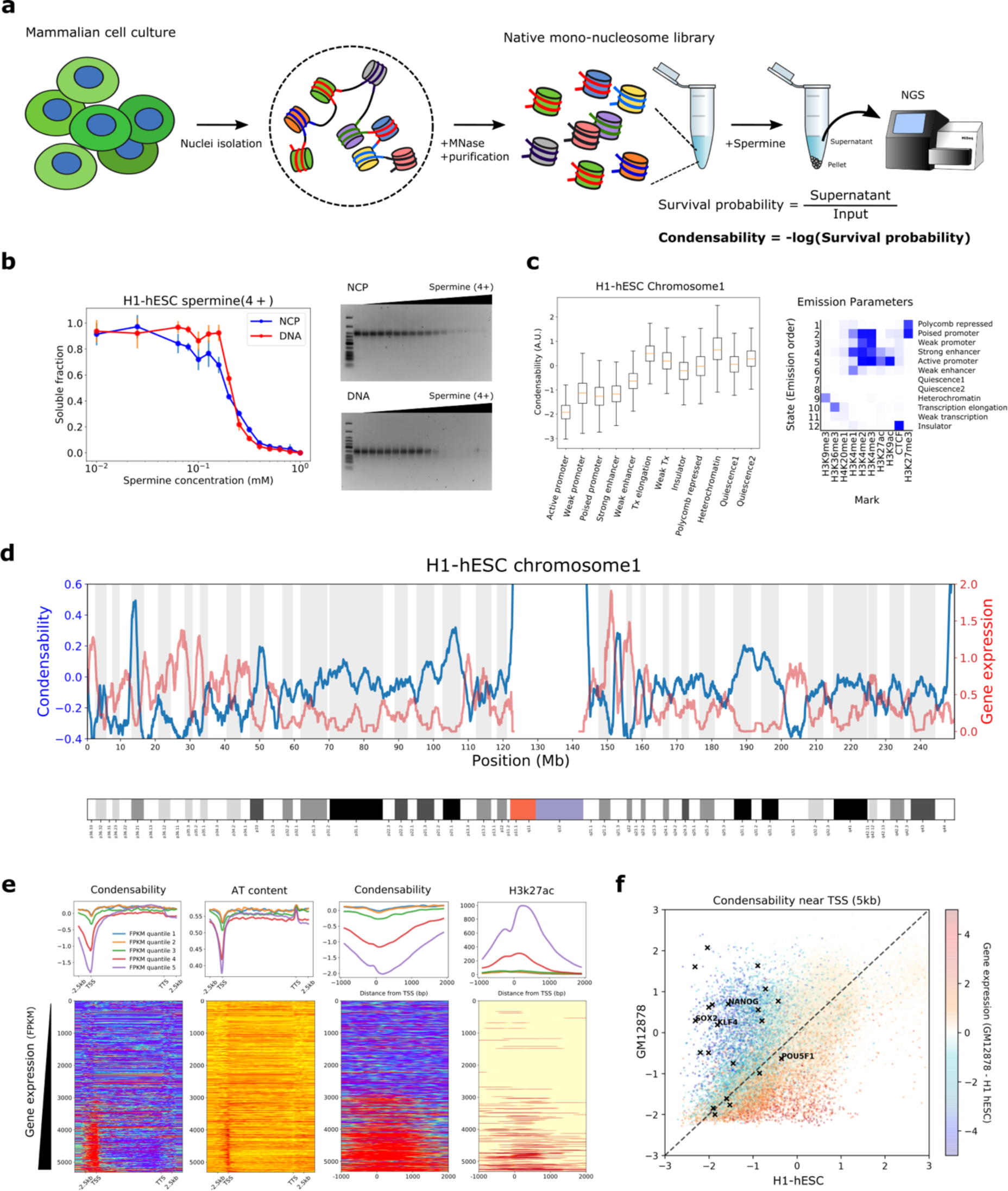
Condense-seq measures single-nucleosome condensability genome-wide. **a**, Overall workflow of condense-seq. **b**, The total amount of nucleosome core particles (NPC) or nucleosomal DNA remaining in the supernatant was measured by UV-VIS spectrometry (left) and their integrity was checked by running gels (right, lane 1 is for DNA NEB Low Molecular Weight ladder) for different concentrations of condensing agents. **c**, Genome segmentation into chromatin states based on histone PTM ChIP-seq data (right). All mono-nucleosomes of chromosome 1 were categorized, and their condensability distribution for each chromatin state is shown. **d**, RNA-seq data (red) and condensability (blue) over the entire chromosome 1 (bin size 100 kb). **e**, All genes in chromosome 1 were grouped into five quantiles according to the transcription level (quantile 1 through 5 for increasing transcription). (top) Condensability, AT content and H3K25ac along the transcription unit coordinate averaged for each quantile. (bottom) Heat maps show the same quantities for each gene, rank ordered with increasing condensability. TSS (transcription start site), TTS (transcription termination site). **f**, Promoter condensability (averaged over 5kb window around TSS) for H1-hESC and GM12878. Each gene is colored according to their relative expression levels in the two cell types. Black symbols are for embryonic stem cell marker genes.

### More highly transcribing genes show lower condensability, especially around the transcriptional start site

Fig. 1d and Extended Data Fig. 3a show a chromosome-wide condensability map (H1 human embryonic stem cell (hESC)). At 1 Mb resolution, condensability varies from –2 to +2 with respect to the average set at zero, and it greatly increases in the sub-telomeric and pericentromeric regions. Gene expression, as reported by RNA-seq ^42^, shows a clear anticorrelation with condensability. At a much finer scale, condensability around the transcription start site (TSS) is the lowest in the most highly expressed genes and vice versa (Fig. 1e). These findings are surprising because they indicate that single native nucleosomes ‘know’ if they are in highly transcribed regions or gene promoters by way of their reduced condensability and vice versa. Other features, such as AT content, CpG methylation density, H3K9ac, H3K27ac, and H3K4me3 also showed dependence on gene expression, but individually were poor predictors of condensability profiles across the promoter region (Fig. 1e and Extended Data Fig 3c,d). For example, although AT content is also the lowest around TSS in genes with the highest expression, its dip is much narrower than the condensability dip (Fig. 1e). In another example, H3K27ac, which, while stronger in highly expressed genes, does not show strong correlation with condensability in either width or rank order (Fig. 1e). Notably, even in highly expressed genes, condensability quickly increases as we examine regions away from the TSS and into the gene body (Fig. 1e).

Next, we used ChromHMM ^43^ to segment the genome into 12 chromatin states based on histone modifications and observed clear differences in condensability depending on the chromatin state (Fig. 1c). Promoters and enhancers show the lowest condensability whereas heterochromatin, gene body, Polycomb repressed, and quiescence state show the highest condensability. In addition, strength dependence was observed: strong promoters and enhancers show lower condensability than weak promoters and enhancers. Overall, transcriptionally active chromatin states show low condensability compared to inactive states, with one exception: the gene body shows high condensability, and this is true even in highly expressed genes, as noted earlier (Fig. 1c).

In the UCSC genome browser ^44^ view of ∼40 kb window in human chromosome 1 (Extended Data Fig. 3b), condensability obtained from H1-hESC shows two major minima of approximately 2kb in width and overlapping with cis-regulatory regions, a promoter and an enhancer. The depth of the minima is approximately 2, indicating that the nucleosomes there are about 7.3 times, *e*^2^, less condensable than the average nucleosomes. Both overlapped with CpG islands and also with Dnase I hypersensitivity peaks which, however, are considerably narrower in width than the condensability dips.

We next tested the possibility that the condensability contrast is primarily driven by AT content ^30^, hence independent of cell type or cellular state, by performing condense-seq for a differentiated cell type, GM12878 (Extended Data Fig. 4). Condensability in the 5 kb region surrounding the TSSs of all annotated genes shows wide variations between the two cell types (Fig. 1f). Importantly, genes with higher expression in the differentiated cell (GM12878) than in the embryonic stem cell (H1 hESC) show lower condensability in the differentiated cell and vice versa. Therefore, condensability of the promoter region is cell type dependent, ruling out the possibility that cell-type independent features such as AT content is the primary determinant of promoter condensability. Notably, embryonic stem cell markers, such as nanog, sox2, and klf4, possess promoter regions much less condensable in the embryonic stem cell than in the differentiated cell (Fig. 1f).

### Native mono-nucleosomes encode A/B compartmentalization information

The chromosome-wide anti-correlation between condensability and gene expression raises the possibility that nucleosome condensability is closely associated with eu-/heterochromatin compartmentalization. Hence, we compared the condensability profile with the A/B compartment score obtained from the H1-hESC Micro-C data ^45^. We observed a clear anti-correlation between the condensability and the A/B compartment score in the chromosome-wide mega-base scale ( Extended Data Fig. 5a) and in the 100 kb scales (Fig. 2d). At a finer scale of TADs and their boundaries that are determined by transacting factors such as cohesins and CTCF ^46^, the correlation between the experimental TAD insulation score and the predicted score based the condensability was understandably poorer (Fig. 2b and Extended Data Fig. 5b). Genomic accessibility measured by ATAC-seq ^47^ also shows a correlation with condensability, in which more accessible/opened genomic regions are less condensable, and less accessible/closed genomic regions are more condensable. However, the correlation was biphasic due to the sparsity of accessible regions revealed by ATAC-seq (Fig. 2e,f). ^46^

**Fig. 2.**
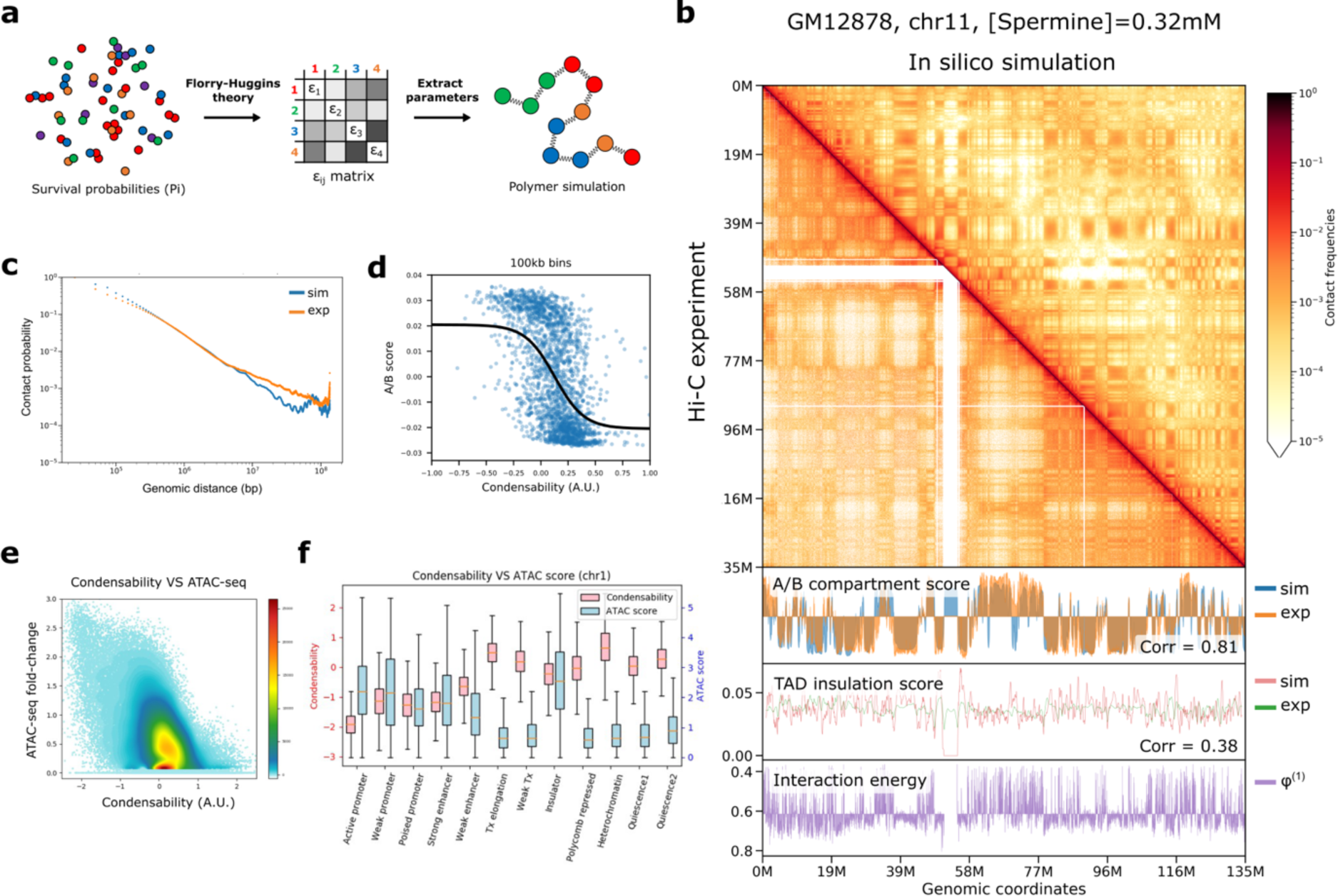
**3D genome compartmentalization information is encoded in native mono-nucleosomes. a**, Nucleosome–nucleosome pair-wise interaction energies, *χ*ij, were derived from the condense-seq measurement based on the Flory–Huggins theory. Chromatin polymer simulation was performed using these interaction energies to predict the three-dimensional chromatin structure solely from nucleosome condensability. **b**, Comparison of contact probability matrix between Hi-C data of GM12878 (lower triangle) and the polymer simulation (upper triangle). In the bottom panel, the A/B compartment scores were computed using the Hi-C data or polymer simulation with interaction energies based on the condensability (phi). TAD insulation scores were also computed for Hi-C data and polymer simulation. **c**, The contact probability vs genomic distance from Hi-C data (orange) and polymer simulation (blue). **d**,A/B compartment score vs condensability in 100 kb bin. Black line is a logistic curve fit. **e**, Condensability vs chromatin accessibility (ATAC score) in 1kb bin. **f**, Condensability and ATAC score vs ChromHMM chromatin state.

We also showed that the *in silico* chromatin polymer simulation of a human chromosome with pair-wise interaction energies derived from the condensability alone as an input (Fig. 2a) can faithfully reproduce A/B compartments from the Hi-C data (correlation coefficient = 0.861 for GM12878) (Fig. 2b,c). This spatial segregation is likely due to the exclusion of less condensable chromatin from the compacted highly condensable core (Extended Data Fig. 5c,d), as demonstrated in the inverted chromatin organization of rod photoreceptors ^48^. Overall, our results imply that the native mono-nucleosomes intrinsically possess, even in the absence of other factors, essentially all the biophysical properties needed for the large-scale A/B compartmentalization of the eukaryote genome.

### Genetic and epigenetic factors collectively determine nucleosome condensability

Next, we sought to identify the genetic and epigenetic features that determine nucleosome condensability. We observed a good correlation between the condensability and the AT content (Extended Data Fig. 6a), reminiscent of stronger polyamine-induced attractive interactions between AT-rich DNA compared to GC-rich DNA of the same length ^30^. No significant correlation was found between condensability and dinucleotide periodicity associated with the rotational phasing of nucleosomal DNA ^29,49^ and extreme DNA cyclizability ^50,51^ (Extended Data Fig. 6b,c), which suggests the independent biophysical mechanisms of nucleosome stability and condensability.

Using DNA methylation and histone ChIP-seq data for H1-hESC in the Encyclopedia of DNA Elements (ENCODE) data portal ^42^, we investigated epigenetic features associated with nucleosome condensability (Extended Data Fig. 6d). Epigenetic marks associated with transcriptional activation are highly enriched in low condensability partitions, with the lone exception of H3K36me3. Repressive epigenetic marks, such as H3K9me3 and CpG methylation density, are more enriched in high condensability partitions. However, some of the other repressive marks, for instance, H3K27me3 and H3K23me2, are enriched in the least condensable fraction (Extended Data Fig. 6d), potentially due to the confounding effects from poised promoters prevalent in embryonic stem cells, which simultaneously possess both active and inactive marks, such as H3K27ac and H3K27me3, respectively ^52–54^ or bivalent promoters in the case of H3K23me2 ^55^. To reduce the confounding effects of diverse features occurring simultaneously in some nucleosomes, we stratified the data into subgroups that shared all features, except one, for comparison with condensability. This conditional correlation analysis showed that high condensability was the most strongly correlated with AT content, H3K36me and H3K9me3 (Fig. 3b). Low condensability was strongly correlated with histone acetylation in general and with H2AFZ, H3K4me1, me2, me3, and H3K79me1, me2. Machine learning based modeling also predicts the nucleosome condensability based on those genetic/epigenetic components as input with similar importance (Extended Data Fig. 6g-I).

**Fig. 3.**
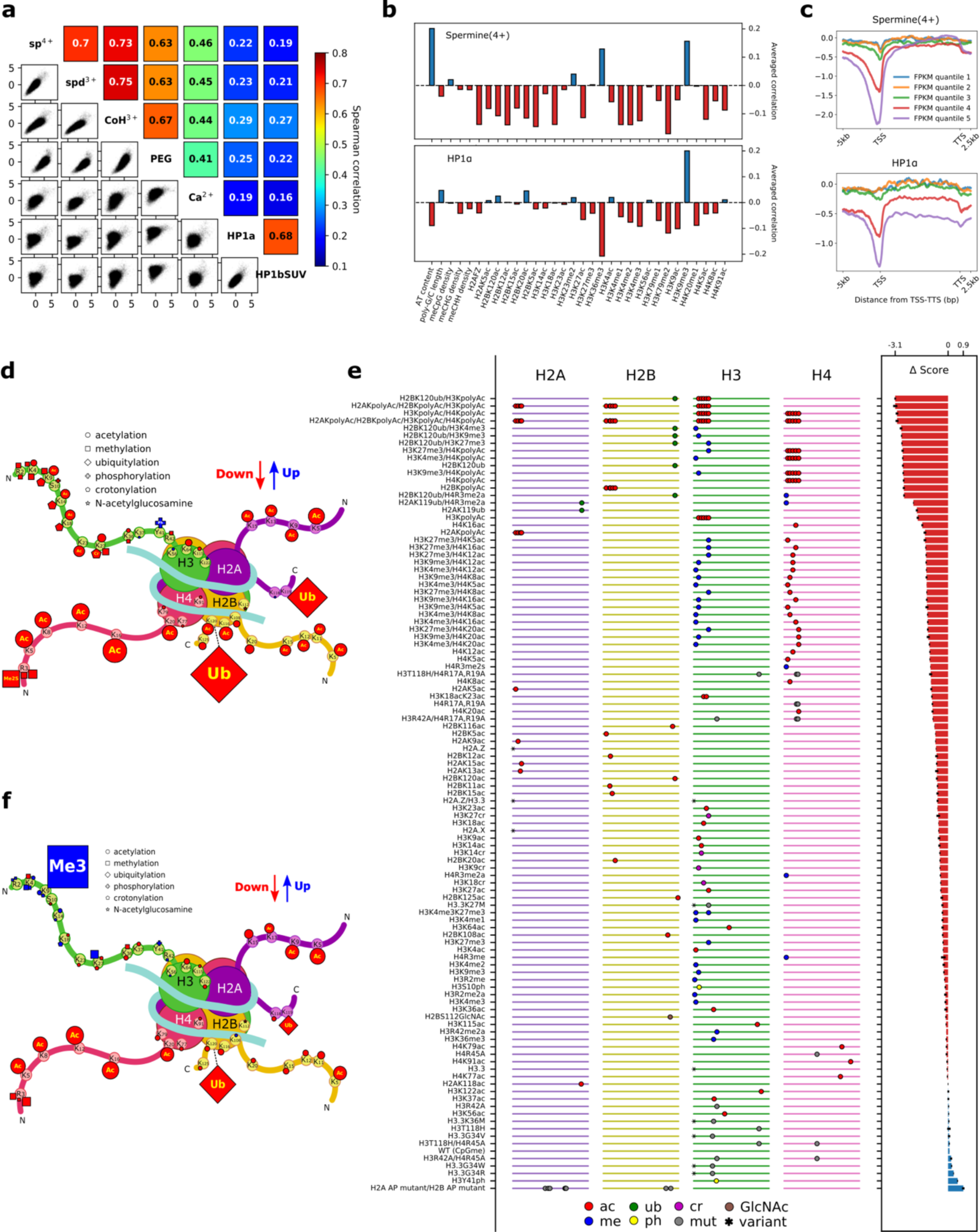
**Identification of the biophysical driving force of chromatin condensation and its genetic/epigenetic determinants. a**, Correlation of condensability scores across condensing agents tested: spermine (sp^4+^), spermidine (spd^3+^), Cobalt hexamine (CoH^3+^), polyethylene glycol molecular weight 8000 (PEG), Ca2+, HP1α and HP1β/tSUV39H1 (HP1bSUV). **b**, Conditional correlations between condensability and various genetic/epigenetic factors for spermine (top) and HP1α (bottom). **c**, Condensability profiles vs gene unit position averaged over each of the five quantiles, from weakly expressed to highly expressed genes for spermine (top) and HP1α (bottom). **d**–**f**, condense-seq results of PTM library. The effects of single PTMs on nucleosome condensation are depicted in the cartoon for spermine (**d**) and HP1α (**f**). Each symbol represents a PTM of a specific type as shown in the legend and its size is proportional to the strength of the effects. The colors of the marks indicate the direction of the effect (red: decrease condensability, blue: increase condensability) compared with the unmodified control. **e**, All condensability scores of the PTM library using spermine as a condensing agent. The library members were sorted based on the lowest to highest condensability scores from top to bottom. On the left panel, the ladder-like lines represent each histone peptide from N-terminal (left) to the C-terminal (right). Each mark on the line indicates the location of the PTMs, and the shape of the marks represents the PTM type (ac: acetylation, me: methylation, cr: crotonylation, ub: ubiquitylation, ph: phosphorylation, GlcNAc: GlcNAcylation, mut: amino acid mutation, var: histone variant). On the right panel,the change in condensability scores of the various modified nucleosomes compared to the control nucleosomes without any PTMs are shown.

We also used bottom-up mass-spectrometry to identify histone PTMs enriched in supernatant/pellet/input native nucleosome samples before and after condensation by spermine (Extended Data Fig. 7). By counting histone H3 and H4 peptides containing PTMs, we computed the enrichment of each PTMs in the supernatant and compared them with unmodified peptides as the control (Extended Data Fig. 7a,b). Consistent with the genomic analysis based on ChIP-seq data, we found that most repressive marks, such as H3K9me3, were mostly depleted, but most of acetylation marks, especially poly-acetylation marks, were strongly enriched in the supernatant. H3K27 and H3K36 methylation marks did not show either clear enrichment or depletion similar to the condense-seq analysis.

To more directly investigate how histone PTMs affect nucleosome condensation without contributions from DNA sequence or cytosine methylation, we used a synthetic nucleosome PTM library formed on identical Widom 601 DNA sequences ^56^. Performing condense-seq and demultiplexing using the appended barcodes, we obtained the condensability change for each PTM mark compared with controls that do not have any PTM marks (Fig. 3e). All single modifications, except for phosphorylation, showed a decrease in condensability relative to the unmodified control (Fig. 3d). Ubiquitylation is the most effective in making nucleosomes less condensable, followed by acetylation, crotonylation, and methylation, in that order. The intrinsic solubilizing effect of ubiquitin-like proteins was previously demonstrated for SUMO ^57^. Electrostatic interaction is a key determinant as shown by the strong impact of acetylation and crotonylation which add negative charges that would requires more positively charged polyamine molecules to neutralize the net negative charged nucleosomes during condensation. Acetylation on histone tails has a much stronger effect than acetylation on the histone fold domain (Fig. 3d), having the strongest effect on the H4 tail, followed by the H2A, H2B, and H3 tails, respectively. In our results, the H2A.Z variant shows significantly reduced condensability compared with the canonical histones (Fig. 3e), which is consistent with the conditional correlation analysis (Fig. 3b) and with previous reports that H2A.Z makes oligonucleosomes more soluble, potentially due to the different acidic patch structure of the variant ^58,59^.

In the cellular context, because genomic nucleosomes are decorated with the combinations of multiple PTMs and cytosine methylation in different sequence contexts as shown in the NMF clustering (Extended Data Fig. 6e,f), nucleosome condensation properties are likely to be a complex emergent outcome of the combined effects of the individual genetic and epigenetic features.

### Electrostatic interaction between nucleosomes is a major biophysical driving force for A/B compartmentalization

Polyamines are thought to induce condensation of DNA and nucleosomes by making ion-bridges between negatively charged DNA ^34,60^. If such charge–charge interaction is a major driving force, other ionic condensing agents should also induce condensation. We performed condense-seq of H1 hESC mono-nucleosomes using spermidine, cobalt-hexamine, magnesium, and calcium ion as well as polyethylene glycol (PEG) (Extended Data Fig. 8a). In all agents, chromosome-wide mega-base scale condensation profiles were anticorrelated with gene expression (Extended Data Fig. 8b). Except for calcium, which poorly condensed mono-nucleosomes, all ionic condensing agents showed good correlations with each other in condensability at the 10 kb scale (Fig. 3a). In addition, all ionic condensing agents also showed very high correlations on the synthetic PTM library (Extended Data Fig.9a-c,f). Intriguingly, the charge conversion mutations on the acidic patch of histone H2A/B, which was previously suggested to be the nucleosome–nucleosome interaction interface ^61–64^, showed the largest condensability changes among the PTM library members for all ionic condensing agents (Fig 3e). Thus, electrostatic interaction between nucleosomes mediated by multivalent ions is probably the main driving force behind large-scale genomic compartmentalization.

Next, we performed the condense-seq of H1 hESC using HP1α and HP1β proteins as condensing agents (Extended Data Fig. 8a). On the mega-base scale, the chromosome-wide condensability profile was anticorrelated with gene expression (Extended Data Fig. 8b) as in the case of ionic agents. However, on the 10 kb scale, condensability did not show good correlations between the ionic agents and the HP1s (Fig. 3a). Using previously annotated data, we quantified the correlations between condensability and various markers of nuclear sub-compartments (LAD: Lamin Associated Domain ^42^, NAD: Nucleolar Associated Domain ^65^, SPAD: SPeckle Associated Domain ^66^) (Extended Data Fig. 8d-f). In all condensing agents, condensability is strongly anticorrelated with nuclear speckle and transcription markers and is weakly anticorrelated with Polycomb markers. Heterochromatin, nucleolar, and lamin-associated marks show a positive correlation with condensability, with the strongest correlation being observed between HP1- mediated condensability and the H3K9me3 marks. Differences between the ionic agents and HP1s were further identified in the ChromHMM genome segmentation: condensability is low at promoters and enhancers for all condensing agents, but the magnitude of this effect was much reduced in HP1 (Extended Data Fig. 4c). Interestingly, the gene body showed low condensability with HP1, in contrast to high condensability with the ionic agents. Consistently, the condensability profile of HP1α from TSS to TTS also shows reduced condensability in highly expressed genes, not only near TSS but also along the gene body (Fig. 3c). Conditional correlations also reveal that condensability with HP1α is negatively correlated with H3K36me3 and positively correlated with H3K9me3 (Fig. 3b).

We also performed condense-seq of the PTM library using HP1α as the condensing agent. H3K9me3 profoundly increases nucleosome condensation by HP1α (Extended Data Fig. 9d, Fig. 3f), which is consistent with HP1α’s role as an H3K9me3 heterochromatin mark reader ^67,68^. Interestingly, regardless of PTM type, most PTMs on the H3 tail also show a slight increase in HP1-induced condensation, and this trend is stronger at locations farther from the nucleosome core. This finding potentially implies that HP1α can weakly recognize other PTMs on the H3 tail in a non-specific manner. Other than H3 tail modifications, most PTMs showed similar effects between HP1α and ionic agents, reducing condensability.

### Polyamine depletion globally hyperpolarizes chromatin condensability but causes local disorganization

Polyamines are one of the most prevalent biological metabolites ^40,41^. We performed condense- seq on mouse T cells, whose activation and differentiation are critically impacted by polyamines ^69^. We isolated and activated CD8^+^ T cells from control mice and mice with a T cell specific deletion in ornithine decarboxylase (ODC) (Fig. 4b), which is a rate limiting enzyme for polyamine synthesis, converting ornithine to putrescine, which can then be further metabolized to spermidine and spermine (Fig. 4a). We also examined wild type mouse CD8^+^ T cells treated with difluoromethylornithine (DFMO), a chemical inhibitor of ODC ^70,71^. In all three (control, ODC KO, and +DFMO), native nucleosomes were purified and subjected to condense-seq with spermine (Fig. 4b and Extended Data Fig. 10a).

**Fig. 4.**
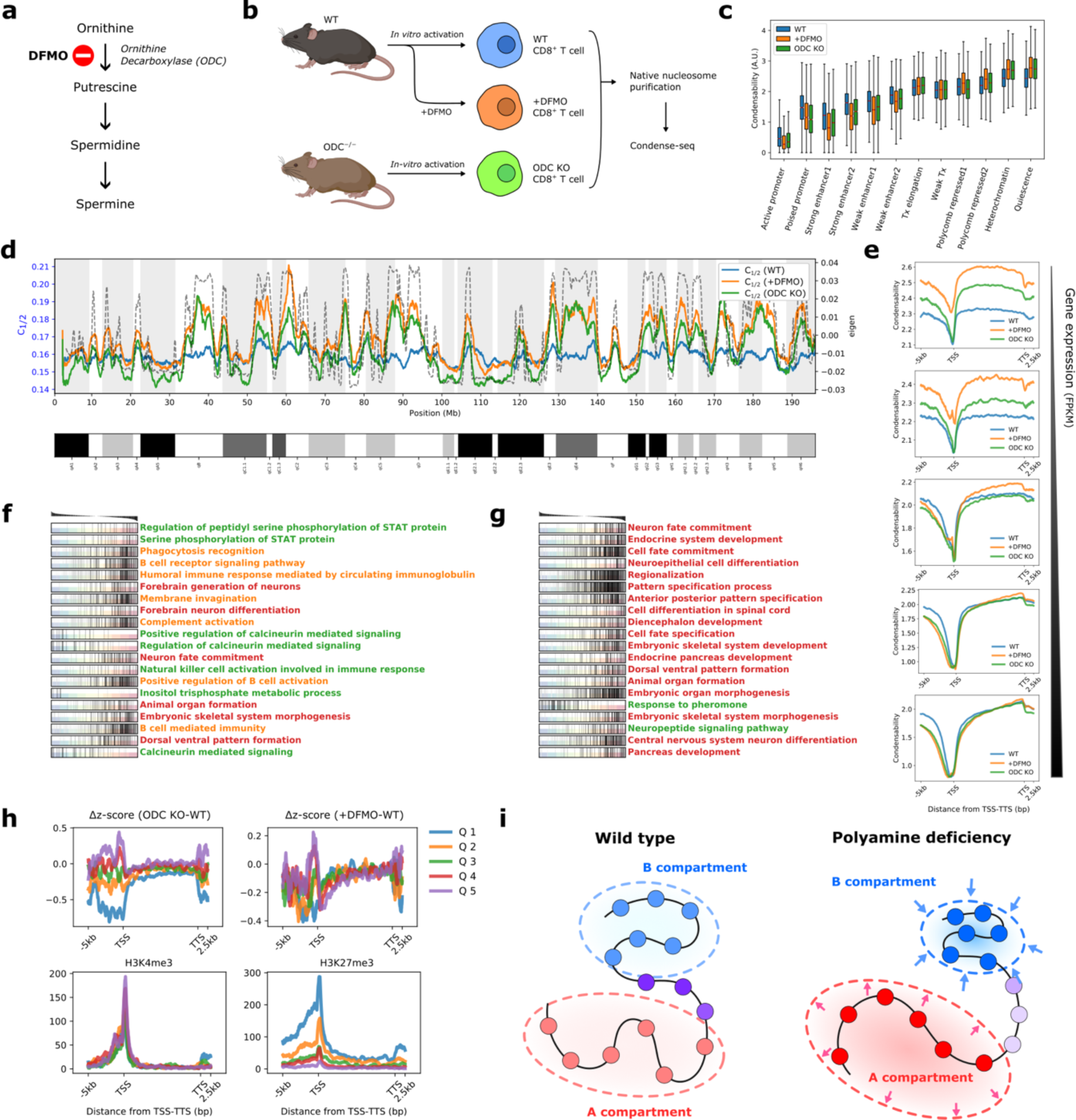
**Polyamine deficiency globally hyperpolarizes but locally disorganizes the chromatin condensability. a**, Ornithine decarboxylase (ODC) is a key enzyme in polyamine biogenesis and is inhibited by DFMO. **b**, Mouse CD8^+^ T cells were activated *in vitro* before condense-seq. wild type (WT), DFMO treated (+DFMO), and ODC knockout (ODC KO). **c**, Mono-nucleosome condensability distribution for WT, +DFMO and ODC KO in various chromatin states classified using ChromHMM. **d**, Condensation point (*c*1/2) for chromosome 1, showing larger dynamic ranger and hyperpolarization for +DFMO and ODC KO. **e**, Condensability over gene units averaged over genes belonging to five quantiles of gene expression (FPKM: Fragments Per Kilobase of transcript per Million mapped reads). **f**,**g**, Gene set enrichment analysis of polyamine deficient conditions (**f**: +DFMO, **g**: ODC KO) compared with the wild type. Genes were ordered by their Δz, z-score of condensability relative to the wild type, shown above. In the graph, each row corresponds to the GO biological process strongly enriched for strongly positive or strongly negative Δz values, and genes belonging to that gene set are localized by tick marks. The enriched GO biological processes are clustered by their biological functions (red: developmental, green: cell signaling, orange: immunologically related). **h**, For each quantile of Δz near TSS, averaged Δz vs transcription unit position is shown for ODC KO VS WT (upper left) and +DFMO VS WT (upper right), and averaged ChIP-seq signals are shown for H3K4me3 (lower left), and H3K27me3 (lower right) **i**, Polyamine deficiency induces global hyperpolarization of chromatin compartmentalization but disrupts local chromatin organization, especially genomic locations enriched with H3K27me3 marks.

For a quantitative analysis of subtle differences across different conditions, we used another metric, condensation point (*c1/2*), a spermine concentration in which the soluble fraction is half the input (Extended Data Fig. 10b). Thus, *c1/2* is inversely correlated with the previously defined condensability score. *c1/2* of nucleosomes has a higher dynamic range in ODC KO and +DFMO compared with wild type (WT) (Extended Data Figure 10c-h), such that disrupting polyamine synthesis appears to amplify the contrast, in which highly condensable nucleosomes become even more condensable and poorly condensable nucleosomes become even less condensable (Fig. 4d). We suggest that when cells cannot rely on polyamines to translate the biophysical properties of nucleosomes into nuclear features such as transcription regulation and chromosome organization, they modify the nucleosomes to accentuate the condensability contrast. In support of this suggestion, similar trends of ‘hyperpolarization’ were observed in individual nucleosomes that were categorized into different chromatin states (Fig. 4c), as well as in the condensability profiles of genes grouped into different quantiles according to their gene expression levels (Fig. 4e).

To investigate the possible local, gene-specific changes upon polyamine depletion, we standardized the condensability score across different conditions using the z-score. ODC inhibition and ODC KO induced z-score changes in single genes, Δz, which are correlated between the two conditions (Pearson’s correlation coefficient 0.62) (Extended Data Fig. 10i). Among the chromatin states, poised promoters and enhancers were the most affected, showing the most significant decrease in condensability upon polyamine depletion (Extended Data Fig. 10j). Gene set enrichment analysis (GSEA) ^72^ showed that many developmental, differentiation, immune signaling, and immune response-related processes are enriched among genes that show significant reduction in condensability upon ODC inhibition (Fig. 4f) or ODC KO (Fig. 4g). Development-related genes, which are repressed through H3K27me3 ^73^, are particularly strongly affected by ODC KO, and indeed, genes with the largest decreases in the promoter condensability upon ODC KO (quintile 1, Fig. 4h) showed the highest enrichment of H3K27me3 (Fig. 4h). Overall, polyamine deficiency not only globally hyperpolarizes genome compartmentalization, making nucleosomes in B compartments and poorly expressed gene promoters more condensable and vice versa, but also causes local chromatin disorganization, especially in developmental genes, which could lead to cell differentiation problems (Fig. 4i).

## DISCUSSION

Our results indicate that biophysical information that is important in large-scale organizations such as A/B compartment and local organizations at the promoter and enhancers is electrostatically encoded within single native nucleosomes. Even when more specific interactions between chromatins and proteins, such as Heterochromatin protein 1 (HP1), Polycomb repressive complex, cohesin and CTCF, and other noncoding RNAs ^74–76^, are responsible for smaller scale, function-directed chromosome organization ^13,38,77,78^, the intrinsic condensability in individual nucleosomes must form a biophysical backdrop that biology must consider (Extended Data Fig. 8g).

The differences in nucleosome condensability between H1-hESC and GM12878 illuminate how compartmentalization changes upon cellular differentiation. The genome-wide condensability in GM12878 shows a higher dynamic range and better correlation with A/B compartment scores (Extended Data Fig. 4b,c). In addition, the condensability near TSS decreased deeply and widely, even affecting toward gene body of highly transcribing genes of GM12878 (Extended Data Fig. 4d), whereas condensability on the gene body of H1-hESC is consistently high regardless of gene expression level (Fig. 1e). This discrepancy could be due to HP1, which polarizes the condensability of genes according to the transcription level in H1-hESC (Fig. 3c).

The PTM library data shows that ubiquitylation, either of repressive H2AK119 or active H2BK120 marks, strongly impedes nucleosome condensation (Fig 3e), suggesting that other factors must be recruited through chemical recognition to differentiate between the two ubiquitin modifications. Interestingly, in the micronuclei where nuclear import is defective, both H2AK119Ub and H2BK120Ub are reduced, potentially contributing to more condensed chromosomes in the micronuclei which are also marked by reduced histone acetylation and increases in H3K36me3 ^79^. We were surprised that almost all PTMs, including charge neutral methylations, reduce condensation. Overall, the direct physical effect of all these modifications is to increase the accessibility of chromatin, albeit to varying degrees depending on their type (Fig. 3d, Extended Data Fig. 9a-c), which might serve as the initial physical opening of chromatin for docking epigenetic readers into action.

Does condensability drive differential gene expression, or is it a mere consequence of differential gene expression? H3K36me3 marks, which are prevalent in highly transcribing gene bodies, do not show an enrichment in low condensability partitions, suggesting that the regions around the TSS such as promoters and enhancers, rather than the gene body itself, are occupied by less condensable nucleosomes. This is further supported by ChromHMM analysis (Fig. 1c) and meta- gene profiles (Fig. 1e). Therefore, high traffic by transcription machinery alone is not sufficient to lower nucleosome condensability, and we favor the model in which cells regulate gene expression through modulating the condensability of promoter nucleosomes. High condensability in the gene body may help avoid spurious transcription initiation.

Polyamines, which exist at ∼ mM concentration in eukaryotic cells ^40^, must play an important role in the process because when they are depleted, cells attempt to compensate by accentuating the contrast in nucleosome condensability (Fig. 4). This hyperpolarization, consistent with the dual role of polyamine as repressors and inducers of gene expression depending on genes and cellular context as previously reported ^80^, can result in various dysfunction.in cell differentiation ^69^, cancer ^81^, and immunology ^82^ via direct interaction or metabolic perturbation of chromatin remodeling, and understanding this link, how polyamines change biophysical properties of chromatin, would be interesting future work to be investigated.

## MATERIAL AND METHODS

### Native mono-nucleosome purification

We used the hydroxyapatite (HAP) based protocol with minor modifications ^39^. See Supplementary Data Note 1 for full details. Briefly, we cultured human embryonic stem cells (H1) and harvested about 100 million cells. Next, we purified the nuclei with 0.3% NP-40 buffer and performed MNase digestion at 37°C for 10 min in the presence of protease inhibitor cocktails and other deacetylation and dephosphorylation inhibitors. The soluble mono-nucleosomes were saved after centrifugation of the insoluble nuclei debris in a cold room. The nucleosome samples were incubated with hydroxyapatite slurry for 10 min, and then unbound proteins were removed by repetitive washing with intermediate salt buffers. Finally, the nucleosomes were eluted with phosphate buffer from the hydroxyapatite slurry. The eluted fraction was checked by extracting DNA from the nucleosome through phenol-chloroform extraction and running a 2% agarose gel. The HAP elution contained mono-nucleosomes, naked DNA and oligonucleosomes. We applied additional size selection of mono-nucleosomes using Mini Prep Cell (Biorad) gel-based size selection purification. The quality of the final mono-nucleosome sample was checked by running a 2% agarose gel and a 20% SDS-PAGE gel. The purified mono-nucleosomes were stored on ice in a cold room for less than a week before the condensation reaction, or they were frozen in liquid nitrogen with 20% glycerol for long-term storage at -80°C.

### Nucleosome condensation assay

The purified native mono-nucleosome sample was extensively dialyzed into 10 mM Tris pH 7.5 buffer through several buffer exchanges using an Amicon Ultra 10kD filter (MilliporeSigma). In each condensation reaction, the final concentration of nucleosome or DNA was 50 ng/µl as DNA weight, and BSA was added to the final 0.2 mg/ml to stabilize the nucleosome core particle. The condensation buffer condition was 10 mM Tris pH 7.5 with additional salt depending on the condensing agents (50 mM NaCl for spermine, and 250 mM NaCl for polyethylene glycol (molecular weight 8000 Dalton)). Eight to 16 samples with different concentrations of condensing agents were prepared simultaneously. They were incubated at room temperature for 10 min and centrifuged at 16,000 g for 10 min, and the supernatant was saved. The soluble nucleosome concentration was measured using a Nanodrop UV spectrometer, and the nucleosome sample integrity was checked by running the 2% agarose gel. The rest of the nucleosomes in the supernatant were saved for use in high throughput sequencing.

### NGS library preparation and sequencing

Using phenol chloroform extraction, genomic DNA was extracted from the nucleosome, which was either the input control sample or the supernatant saved from the nucleosome condensation assay. The extracted DNA sample was then washed several times with distilled water using an Amicon Ultra 10kD filter (MilliporeSigma). Using the NEBNext Ultra II DNA library prep kit (NEB), the DNA was adaptor-ligated and indexed for Illumina NGS sequencing. The final indexing PCR was conducted in 5 to 7 cycles. We used HiSeq 2500 or the NovaSeq 6000 platform (Illumina) for 50bp-by-50bp pair-end sequencing. In each experimental condition, We sequenced the samples over multiple titration points to get 10 kb resolution data but deeply sequenced a few selected titration points to achieved approximately 20x coverage of the entire human genome at the single nucleosome resolution. In this paper, we mainly focused on the titration points near complete depletion of solution fraction, in where we could observe the highest contrast of nucleosome condensabilities with strong selection power (e.g. [spermine]=0.79 mM in Fig. 1b and [HP1α] =6.25 µM in Extended Data Fig. 8a).

### Genetic and epigenetic datasets

All genome references and epigenetic data, including DNA methylation, histone ChIP-seq, and Hi-C, used in this work are shown in Supplementary Table 1-11.

### Computation of genome-wide nucleosome condensability

First, we obtained coverage profiles along the genome for input control, and for the supernatant sample of each titration after the alignment of pair-end reads on the hg38 human genome assembly using the Bowtie2 software ^83^. Based on the coverage profile of the input control data, the position of each mono-nucleosome was localized by calling the peaks or finding the local maxima of the coverage profile. Beginning by randomly choosing a peak, the algorithm searched for all peaks in both directions, not allowing more than 40bp overlaps between 147 bp peak windows. For each nucleosome peak, the area of coverage within a window (we picked 171 bp as the window size) was computed for both the control and supernatant samples. Then, the negative natural log of the ratio of supernatant versus control area was used as a condensability metric for each mono-nucleosome peak. For finer regular sampling used in plotting metagene profiling, the genome was binned into a 167 bp window with 25 bp sliding steps to compute the coverage area and the condensability scores. For a large scale, we binned the genome into 1 kb or 10 kb and counted the reads aligned onto each bin to compute the condensability scores as the negative natural log of the ratio of supernatant to input read counts.

### Computation of a condensation point, *c*1/2

The condensation point, *c*1/2 was computed by using the survival probabilities of nucleosomes in multiple spermine concentrations. For each 10kb genomic bin, we estimated the nucleosome counts in the input and supernatants after condensation in different spermine concentrations. We obtained the data points of spermine concentrations versus the soluble fraction of nucleosomes and fitted them with a logistic function. We then defined C1/2 as the spermine concentration when the soluble fraction is half the input.

### Z-score computations as enrichment metric

We used the z-score as the enrichment metric for genetic and epigenetic features. For example, we counted the number of CpG dinucleotides in each mono-nucleosome and standardized their distribution by subtracting the mean across all nucleosomes and dividing it by the standard deviation. Thus, each mono-nucleosome was assigned with a z-score of the CpG dinucleotide counts as the metric of how enriched or depleted the CpG was compared with the average in the unit of standard deviation. For the partitioned or grouped data set of the quantile analysis, we used the averaged z-score for each partition as the enrichment metric.

### Data stratification and conditional correlation

To minimize the confounding effects between the genetic and epigenetic features of nucleosome condensation, the data were divided into subgroups that had one varying test variable, but all other variables were constant. For example, to evaluate whether the AT-content was correlated with condensability, the data were divided into smaller groups with the same genetic and epigenetic features, such as H3K4me3 and CpG methylations, etc. except for AT content. In each stratified subgroup, we checked the correlation between AT content and condensability. We then defined the conditional correlation between AT content and condensability as the weighted average of all correlations over the stratified subgroups, weighted according to the data size of each subgroup. In practice, it is difficult to obtain enough data for each stratified subgroup when the feature set is high dimensional. In this case, we discretized each genetic–epigenetic feature into a specific number. All histone ChIP-seq scores were discretized into 10 numbers, and other scores were discretized into 100 numbers.

### NMF decomposition

The genetic–epigenetic features of all mono-nucleosomes in chromosome 1 are linearly decomposed into 10 basis property classes through a Scikit-learn NMF Python package. The nucleosomes were clustered into each property class, with the highest component value in linear decomposition.

### Machine learning models

First, we randomly selected 0.1 million nucleosomes from chromosome 1 for machine learning. With this data set, the Ridge regressor, Supported Vector regressor, Gradient Bosting regressor, Random Forest regressor, and multi-layer perception regressor are trained and validated through 10-fold cross-validations. All machine learning training and predictions were performed using the Scikit-learn Python package.

### Wild type human nucleosome reconstitution

Individual human histones, H2A, H2B, H3.1 and H4, were purchased from The Histone Source (Colorado State University) and the octamers were reconstituted and purified following the standard protocol ^84^. Then nucleosomes were reconstituted with Widom 601 DNA by following the standard gradient salt dialysis protocol ^85^. Nucleosomes are further purified by Mini Prep Cell (Bio-Rad) to eliminate naked DNA or other byproduct contaminants. The background Widom 601 DNA was designed to have the same length and sequence as in the PTM library. However, it has different primer-binding sequences, so that it cannot be amplified along with the library members.

### HP1α and HP1β/tSUV39H1 complex purification

We expressed and purified HP1α following the previous protocol ^12^. As a summary, we expressed HP1α with His6 affinity tag in *E. coli* Rosetta (DE3) strains (MilliporeSigma) at 18 °C overnight. After cell lysis, the protein was first purified by cobalt-NTA affinity purification. Then, his-tag was cleaved by TEV protease, which was removed by anion-exchange purification using HiTrap Q HP column (GE Healthcare). HP1α was further purified through size-selection using a Superdex-75 16/60 size-exclusion column (GE Healthcare). Heterochromatin Protein 1 beta with truncated SUV39H1 complex (HP1Δ/tSUV39H1) is similarly purified based on the previous protocol ^38^.

### Nucleosome condensation assay of the PTM library

The PTM library was prepared as previously described ^56^. The nucleosome condensation reaction of the PTM library was performed similarly, as described for the native mono-nucleosomes. However, because of the limited amount of PTM library sample, we spiked only 1% (v/v) sample amount of the library into 99% (v/v) of reconstituted human nucleosomes as background for the condensation reaction. For condensation experiments with HP1α, a final concentration of 50 ng/µl of DNA or nucleosome (DNA weight) was used in the reaction buffer (10 mM Tris-HCl pH 7.5, 100 mM NaCl, 0.2 mg/ml BSA) in the presence of a final 5% (v/v) polyethylene glycol (PEG) 8000 as crowding agent. Various amounts of HP1α were added to start condensation.

### NGS library preparation and sequencing of the PTM library

The DNA sample was purified through phenol-chloroform extraction, followed by several washes with distilled water using the Amicon Ultra filter (MilliporeSigma). The DNA library was then prepared for Illumina NGS sequencing by PCR using Phusion HF master mix (NEB) and custom indexed primers for the PTM library ^56^. During amplification, the background nucleosome DNA is not amplified, as it has different primer-binding sequences. We used MiSeq (Illumina) for sequencing libraries with custom primers, following previous studies ^56^.

### Condensability calculation for the PTM library

The PTM library is demultiplexed based on the DNA hexamer barcodes by using custom Python script and Bowtie2 aligner ^83^. Then, we approximated the nucleosome counts using total soluble fraction information, which was measured by the Nanodrop UV-VIS spectrometer, and the fraction of the individual members in the library, which was measured by Illumina sequencing. Finally, we computed the survival probability of each member in the library, which is the number of the remaining nucleosomes in the solution after condensation over input control. A negative log of survival probability was used for the condensability metric. For the PTM library, condensability averaged over many titration points was used as a condensability score for further analysis.

### Nucleosome**–**nucleosome interaction energy calculations and chromatin polymer simulation

Coarse-grained molecular dynamics simulations of chromatin were carried out using OpenMM software ^86^. Chromatin was modeled as beads-on-a-string polymers with each bead representing a 25-kilobase-long genomic segment. Energy terms for bonds, excluded volume, spherical confinement, and sequence-dependent contacts were defined. Sequence-dependent contact energies were parameterized using read counts from condense-seq experiments. Contact probability matrixes were computed from these simulation trajectories and compared with experimental Hi-C contact maps. Full simulation details are provided in the supplementary information.

### Mouse CD8^+^ T cell culture and in vitro activation

Wild type C57BL/6 mice and mice expressing Cre recombinase (CD4Cre) under the control of the CD4 promoter were purchased from Jackson Laboratories. Odc^flox/flox^ mice were purchased from the KOMP repository. All mice were bred and maintained under specific pathogen free conditions under protocols approved by the Animal Care and Use Committee of the Johns Hopkins University, Baltimore, USA, in accordance with the Guide for the Care and Use of Animals. Mice used for all experiments were littermates and matched for age and sex (both male and female mice were used). Mice for all strains were typically 8-12 weeks of age.

Naïve CD8^+^ T cells were isolated from the spleen 8- to 12-wk-old mice using a negative selection CD8 T cell kit (MojoSort Mouse CD8 T Cell Isolation Kit) according to the manufactureŕs protocol. Isolated T cells (1 × 106/mL) were activated using plate bound α-CD3 (5 μg/mL) and soluble α- CD28 (0.5 μg/mL) in T cell media (TCM; 1640 Roswell Park Memorial Institute [RPMI] medium with 10% fatal calf serum, 4 mM L-glutamine, 1% penicillin/streptomycin, and 55 μM beta-mercaptoethanol) supplemented with 100 U/mL rhIL-2 (Peprotech). Cells were cultured at 37°C in humidified incubators with 5% CO2 and atmospheric oxygen for 24 h following activation. After 48 hours T cells were removed from α-CD3 and α-CD28 and cultured at a density of (1x106/ml) in rhIL-2 (100 U/mL) at 37 °C for 7 days, with a change of media and fresh rhIL-2 every 24 hours.

To inhibit ornithine decarboxylase (ODC), cells were incubated with difluoromethylornithine (DFMO) 2.5mM for 24 hours at day 6 of culture. ODC ^-/-^, Wildtype and DFMO treated cells were harvested at day 7 for chromatin isolation and sequencing.

### Histone PTM enrichment measurement using bottom-up mass-spectrometry

For the mass spectrometry measurement, the native mono-nucleosome was purified from the GM12878 cell line, and a nucleosome condensation assay was similarly performed using spermine (250 ng/µl nucleosome, 0.079 mM spermine in 10m M Tris-HCl pH 7.5 buffer at room temperature). The input/soluble/pellet nucleosome sample was washed several times in 10 mM Tris-HCl pH 7.5 buffer using an Amicon Ultra filter (10 kD cutoff) to remove spermine and kept at 70 °C for 20 min to dissociate DNA from the histones. The free DNA was further removed in the desalting step of the mass spectrometry process. About 20 µg of purified histones was derivatized using propionic anhydride ^87^ followed by digestion with 1 µg trypsin for bottom-up mass spectrometry. The desalted peptides were then separated in a Thermo Scientific Acclaim PepMap 100 C18 HPLC Column (250mm length, 0.075mm I.D., Reversed Phase, 3 µm particle size) fitted on an Vanquish™ Neo UHPLC System (Thermo Scientific, San Jose, CA, USA) using the HPLC gradient as follows: 2% to 35% solvent B (A = 0.1% formic acid; B = 95% MeCN, 0.1% formic acid) over 50 minutes, to 99% solvent B in 10 minutes, all at a flow-rate of 300 nL/min. About 5 µl of a 1 µg/µl sample was injected into a QExactive-Orbitrap mass spectrometer (Thermo Scientific) and a data-independent acquisition (DIA) was carried on, as described previously ^87^ Briefly, full scan MS (m/z 295−1100) was acquired in the Orbitrap with a resolution of 70,000 and an AGC target of 1x106. Tandem MS was set in centroid mode in the ion trap using sequential isolation windows of 24 m/z with an AGC target of 2x105, a CID collision energy of 30 and a maximum injection time of 50 msec. The raw data were analyzed using the in-house software, EpiProfile ^88^. The chromatographic profile and isobaric forms of peptides were determined using precursor and fragment extracted ions. The data were output as peptide relative ratios (%) of the total area under the extracted ion chromatogram of a particular peptide form to the sum of unmodified and modified forms belonging to the same peptide with the same amino acid sequence. The log2 fold change in the peptide relative ratio in the soluble/pellet fraction versus the input was computed as the enrichment metric. Using the unmodified peptide as the reference, the difference in fold change (delta fold change) between the PTM modified peptide and the unmodified peptide was computed and plotted as a heatmap.

## DATA AVAILABILITY

Sequencing data were deposited in the GEO database with accession number GSEXXXXXX

## CODE AVAILABILITY

All condense-seq data analysis was conducted using custom Python scripts which can be found on https://github.com/spark159/condense-seq.

## SUPPLEMENTARY DATA

Supplementary Data are available online.

## Supporting information

park_extended

## ACKNOWLEDGMENTS

We thank Jejoong Yoo for formulating the initial project idea and Kotaro Onishi and Andrew Feinberg for experimental advice. We also thank Winston Timp, Gaku Mizuguchi and Daehwan Kim for their helpful discussions. The HP1α plasmid is a gift from the Geeta Narlikar lab at UCSF. HP1β and SUV39H1 plasmids are gifts from the Pilong Li lab at Tsinghua University.

## CONTRIBUTIONS

S.P. and T.H. designed the research. S.P. performed all aspects of the research and data analysis. S.P. and T.H. wrote the paper. Other authors contributed to the following areas: A.A. performed interaction energy calculations and chromatin polymer simulations. G.E.C. maintained mouse lines and purified/cultured CD8^+^ T cells. N.A.B. helped the cell culture and nucleosome purification. M.M.M. advised on PTM library-based experiments. N.V.B. performed bottom-up mass spectroscopy for identifying histone PTM marks. B.Z., B.A.G., T.W.M., and E.L. P. provided helpful scientific discussion and supported scientific collaboration. All authors commented on the manuscript.

