## Supplementary material for "Electrostatic encoding of genome organization principles within single native nucleosomes": park_extended

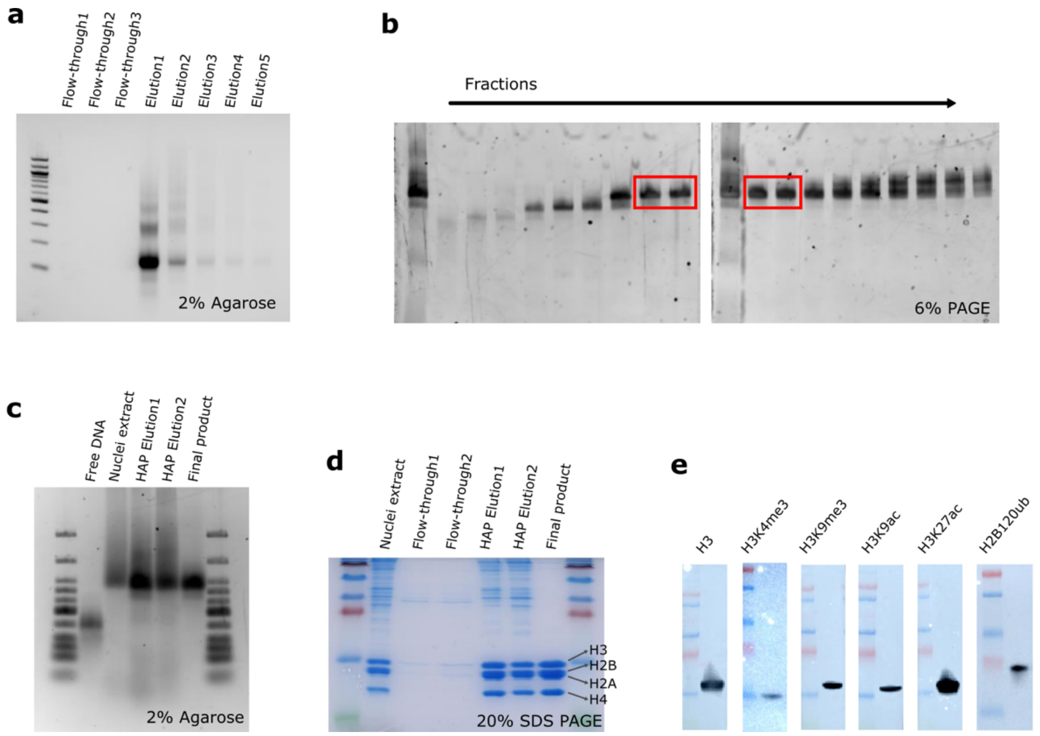


**Extended Data Fig. 1: Intact native mono-nucleosomes obtained by hydroxyapatite (HAP) and size-selective purification.**

**a**, After the HAP purification of MNase-treated chromatin, flow-through and elution samples were run in 2% agarose gel. **b**, Mono-nucleosomes were selected through further size-selective purification of HAP elution. Each purification step and the quality of final product was validated by running the samples in 2% agarose gel (**c**), SDS-PAGE gel showing only four histones without other proteins (**d**), and western blot for histone PTMs (**e**).

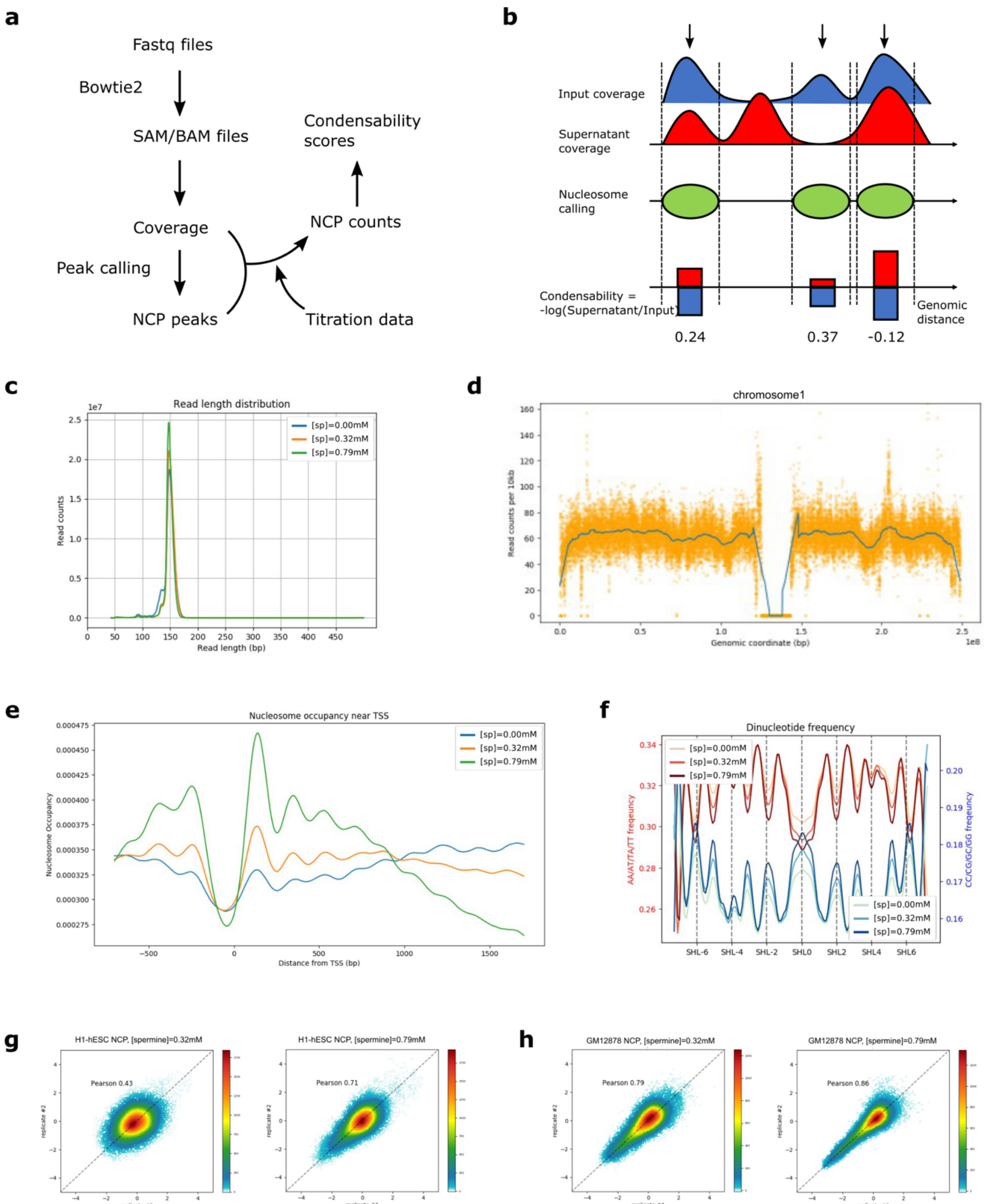


**Extended Data Fig. 2: Computational pipeline and data quality controls for condense-seq.**

**a**,**b**, The pipeline of Condense-seq analysis is composed of (i) reads alignment by Bowtie2, (ii) coverage calculations, (iii) mono-nucleosome peak calling for each local maximum of input coverage, (iv) absolute nucleosome count estimation using coverage area and soluble fraction changes, and (v) compute condensability score as minus log of soluble fraction after condensation for each nucleosome. For the quality control of mono-nucleosome samples, we checked that the length distribution of nucleosomal DNA is mostly at 150 bp (**c**), reads are covered evenly along the genome (**d**), nucleosome positioning shows the signature properties such as the nucleosome free region (NFR) near the transcription start site (**e**), and the periodicity of AT-rich versus GC-rich dinucleotides in nucleosomal DNAs (**f**). Correlation between replicates of Condense-seq measurements on H1-hESC (**g**) and GM12878 (**h**) are shown for condensabilities determined for 0.32 mM spermine and 0.79 mM spermine.

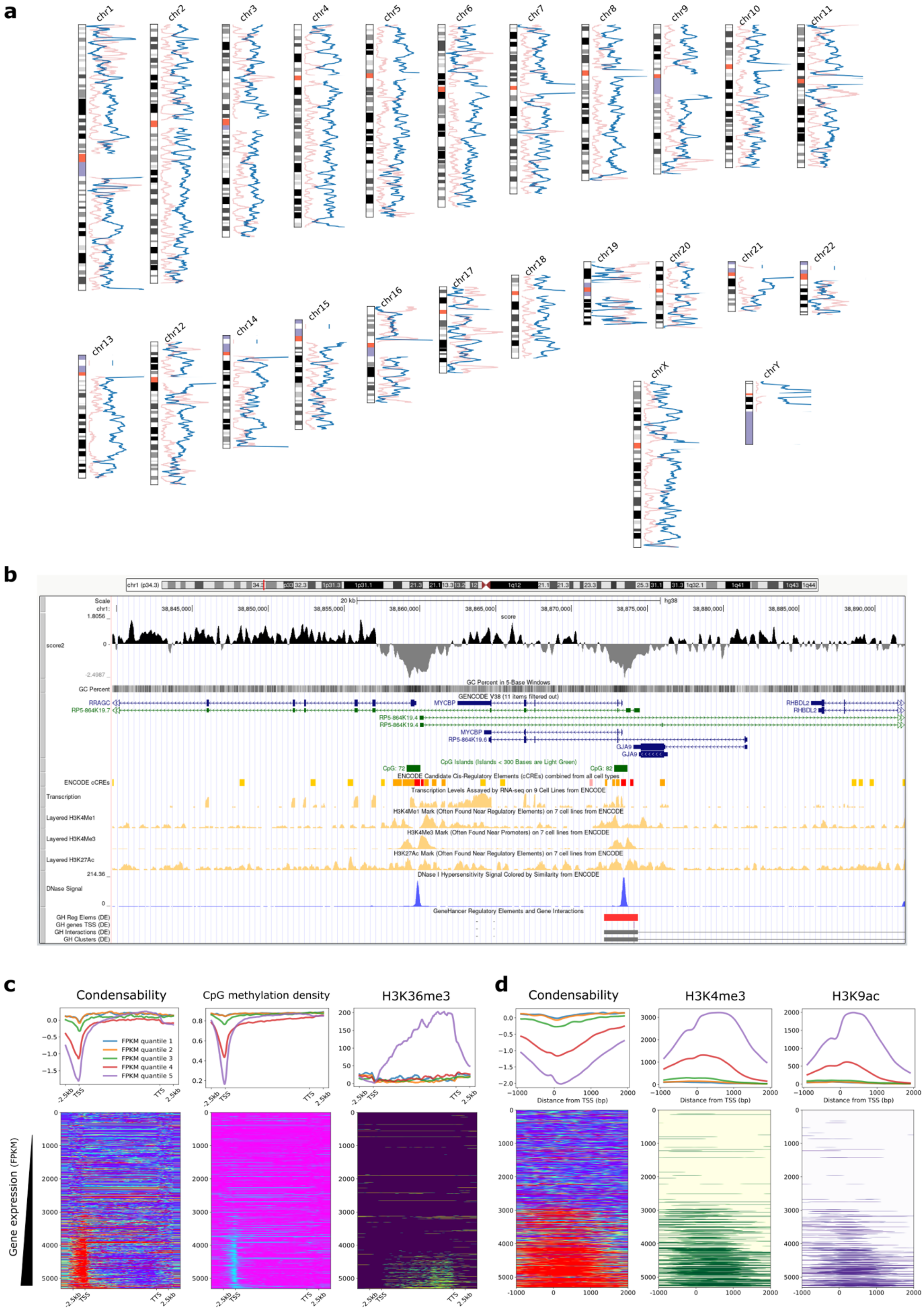
a
b
c
d
Condensability
CpG methylation density
H3K36me3
H3K4me3
H3K9ac
FPKM quantile 1
FPKM quantile 2
FPKM quantile 3
FPKM quantile 4
FPKM quantile 5
Distance from TSS (bp)
Gene expression (FPKM)
GENCODE V38
GC Percent
ENCODE cCREs
DNase Signal

**Extended Data Fig. 3: Condensability scores of H1-hESC across all chromosomes and correlations with gene expression and other epigenetic marks near the TSS.**

**a**, Comparison between condensability (blue) and transcription level (red) along all chromosomes. **b**, Snapshot of UCSC genome browser for the condensability profile along with many other cis-regulatory elements. **c**,**d**, All genes in chromosome 1 were grouped into five quantiles according to the transcription level (quantile 1 through 5 for increasing transcription). (top) Condensability, meCpG fraction, H3K36me3, H3K4me4 and H3K9ac along the transcription unit coordinate averaged for each quantile. (bottom) Heat maps show the same quantities for each gene, rank ordered with increasing condensability. TSS (transcription start site), TTS (transcription termination site). Views zoomed around TSS are shown in (**d**) .

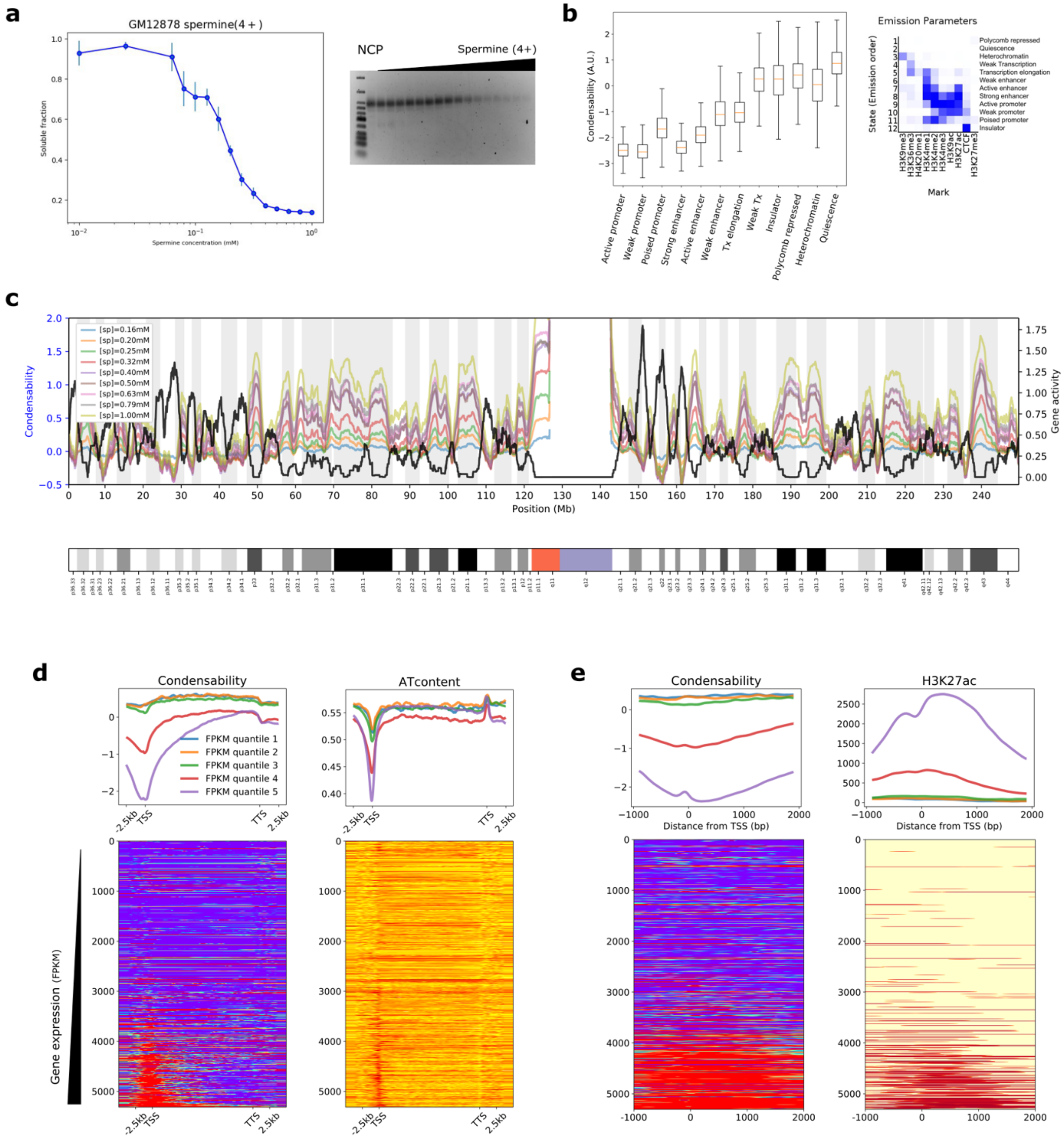


**Extended Data Fig. 4: Condense-seq measurement of native mono-nucleosomes from GM12878 cells and the comparison between nucleosome condensability and their chromatin states.**

**a**, Native mono-nucleosomes were purified from GM12878 cell line, and the condensation was induced by adding spermine. Soluble fractions were measured using UV-VIS spectroscopy (left) and ran in the 2% agarose gel (right). **b**, Genomic nucleosomes were grouped according to their ChromHMM chromatin states based on the combination of various PTMs. Their condensabilities are shown in box plot for each chromatin state. **c**, The condensability score was plotted over human chromosome1 for different spermine titration points (colored lines) and compared with the gene expression level (black line). The condensability profiles from TSS to TTS (**d**) near TSS (**e**) for five quantiles based on the gene expression levels in the GM12878 cell line. Also shown at AT content and H3K27ac profiles.

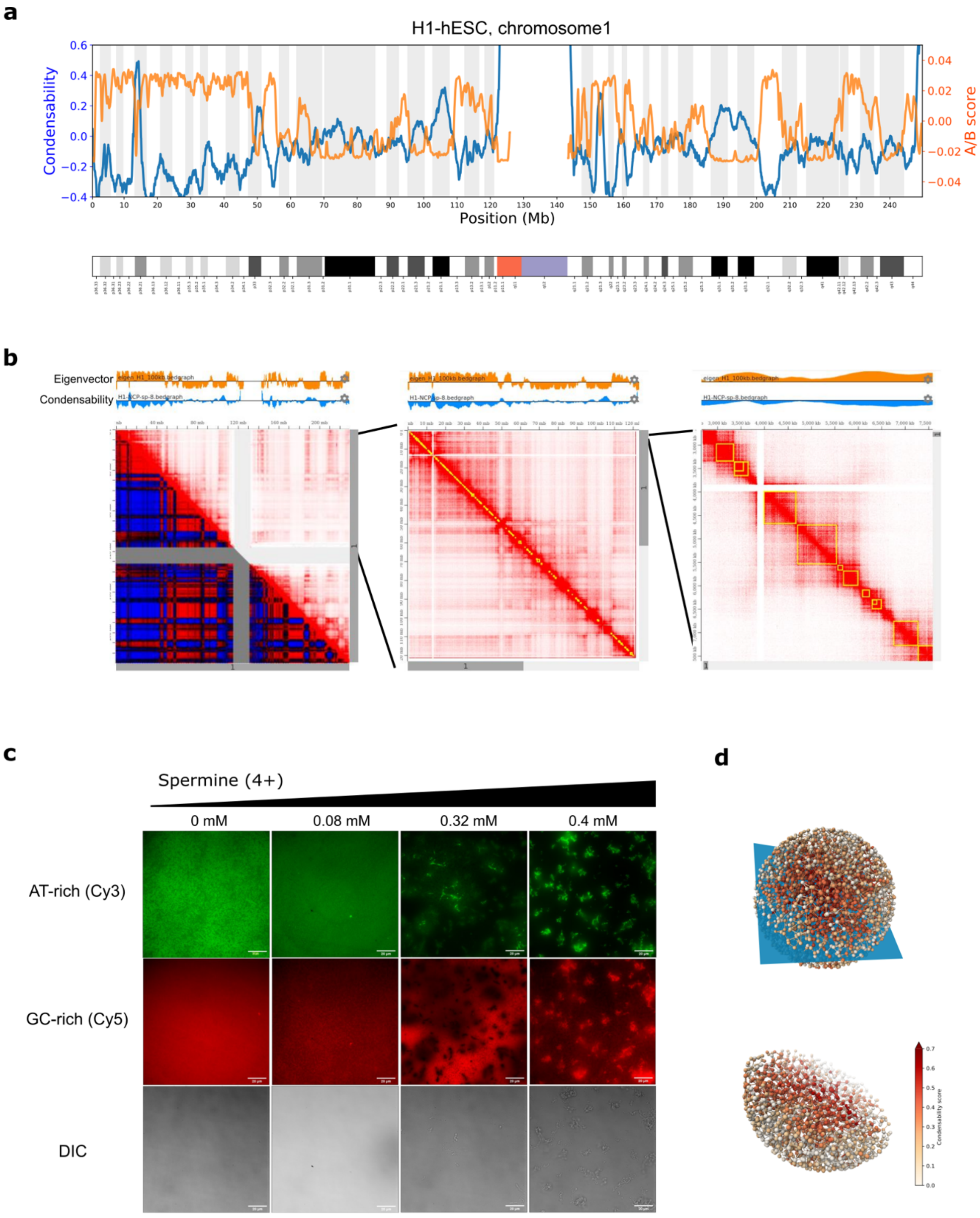


**Extended Data Fig. 5: Spatial separation of molecules promoted by condensability difference to compartmentalize the genome.**

H1-hESC condensability (blue) and A/B compartment scores based on Micro-C data (orange) in mega base-pair resolution of chromosome 1 (**a**) and finer resolution (**b**). **c**, PCR amplified AT-rich (Cy3 labeled) and GC-rich (Cy5 labeled) DNAs were equally mixed and condensed in various spermine concentrations. For each condition, DNA condensates were imaged using wide-field microscope. As spermine concentration increased, AT-rich DNAs formed a condensed core first, and GC-rich DNAs condensed over the AT-rich core at higher spermine concentrations, promoting the spatial separation between AT-rich versus GC-rich condensate. **d**, Chromosome polymer simulation with condense-seq data using spermine as only input (GM12878, chr12) shows that highly condensable chromatin is compacted into the core and the rest is excluded to generate spatially separate compartments (red = condensability score).

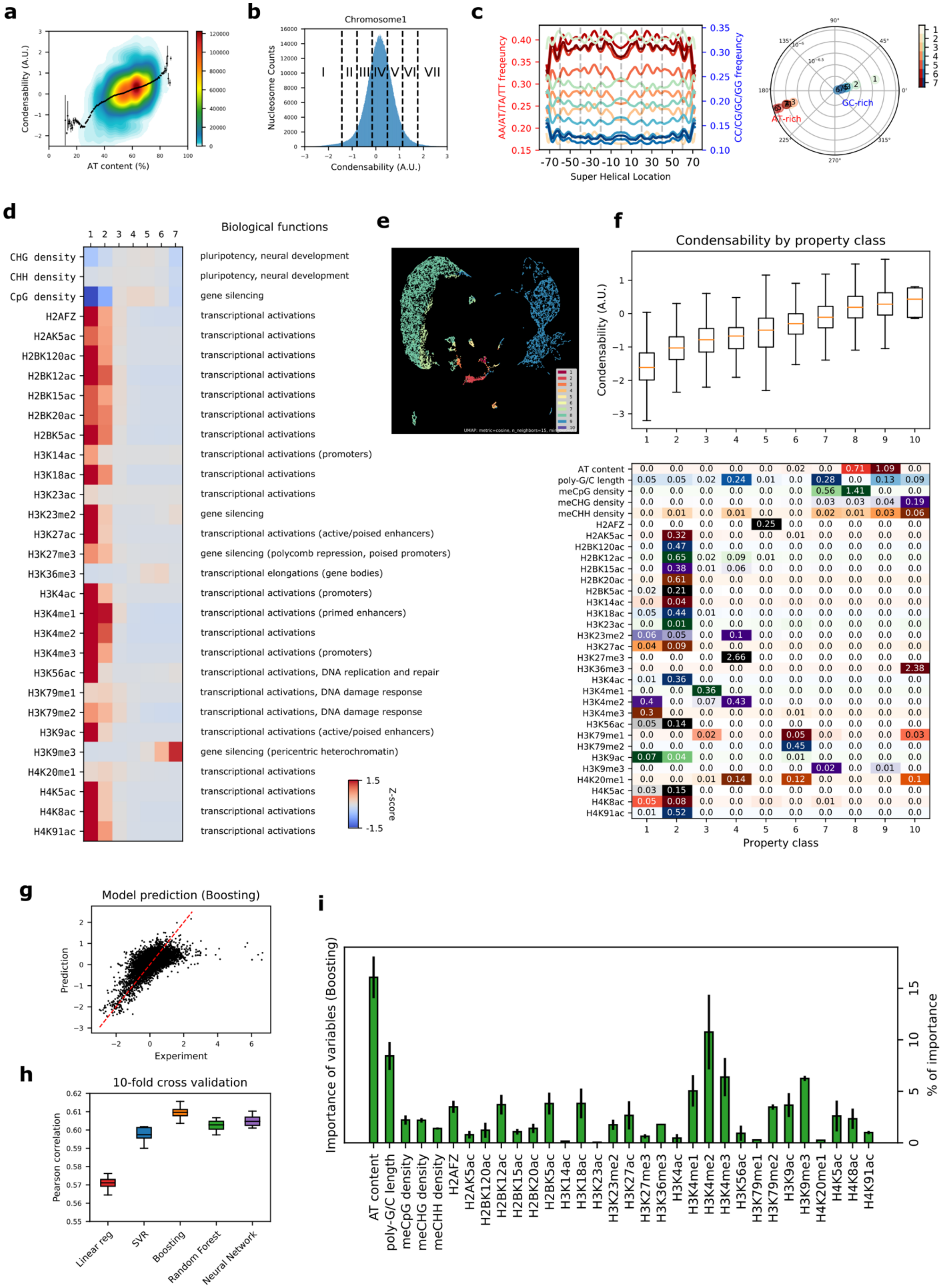
a
AT content (%)
Condensability (A.U.)
b
Chromosome1
Nucleosome Counts
Condensability (A.U.)
I II III IV V VI VII
c
AA/AT/TA/TT freqeuncy
CC/CG/GC/GG freqeuncy
Super Helical Location
AT-rich
GC-rich
d
Biological functions
Z-score
e
f
Condensability by property class
Condensability (A.U.)
Property class
g
Model prediction (Boosting)
Prediction
Experiment
h
10-fold cross validation
Pearson correlation
Linear reg
SVR
Boosting
Random Forest
Neural Network
i
Importance of variables (Boosting)
% of importance

**Extended Data Fig. 6: Deciphering the genetic and epigenetic determinants of genomic nucleosome condensation.**

**a**, Scatter plot of the condensability of mono-nucleosomes in chromosome 1 and the AT contents of corresponding nucleosomal DNA. **b**, The nucleosome population was partitioned into seven partitions, from low to high condensability. The periodicity of AT-rich versus GC-rich dinucleotides was analyzed. The results for each partition (**c**, left), and its amplitude and phase were computed using Fourier transformation and represented in polar plot (**c**, right, radius: amplitude, angle: phase). **d**, The enrichment analysis of all DNA methylation and histone ChIP-seq data available in ENCODE over different condensability partitions from low to high (1-7 partitions in **b**). **e**, After randomly selecting 1% of the mono-nucleosomes in chromosome 1, their genetic and epigenetic features were projected into two-dimensional scatter plots through UMAP embedding. Each color represents one of the property classes of the NMF analysis shown in panel **f**. **f**, The genetic and epigenetic features of all mono-nucleosomes in chromosome 1 were linearly decomposed into 10 property classes by NMF algorithms. Each property class has a specific combination of features, as shown in the matrix (lower panel). All nucleosomes were clustered into each representative property class with the biggest contributions. After clustering, nucleosome condensabilities were plotted as boxplot for each class (upper panel). **g–l**, Multivariate linear regression (linear reg), Supported Vector Machine regression (SVM), gradient boosting regression (Boosting), random forest regression (Random Forest), and neural networks were used to predict nucleosome condensability. All showed similar correlations between experimental values and predictions in a 10-fold cross-validation (**g**,**h**). The importance of genetic–epigenetic features in prediction was computed using the boosting method (**i**).

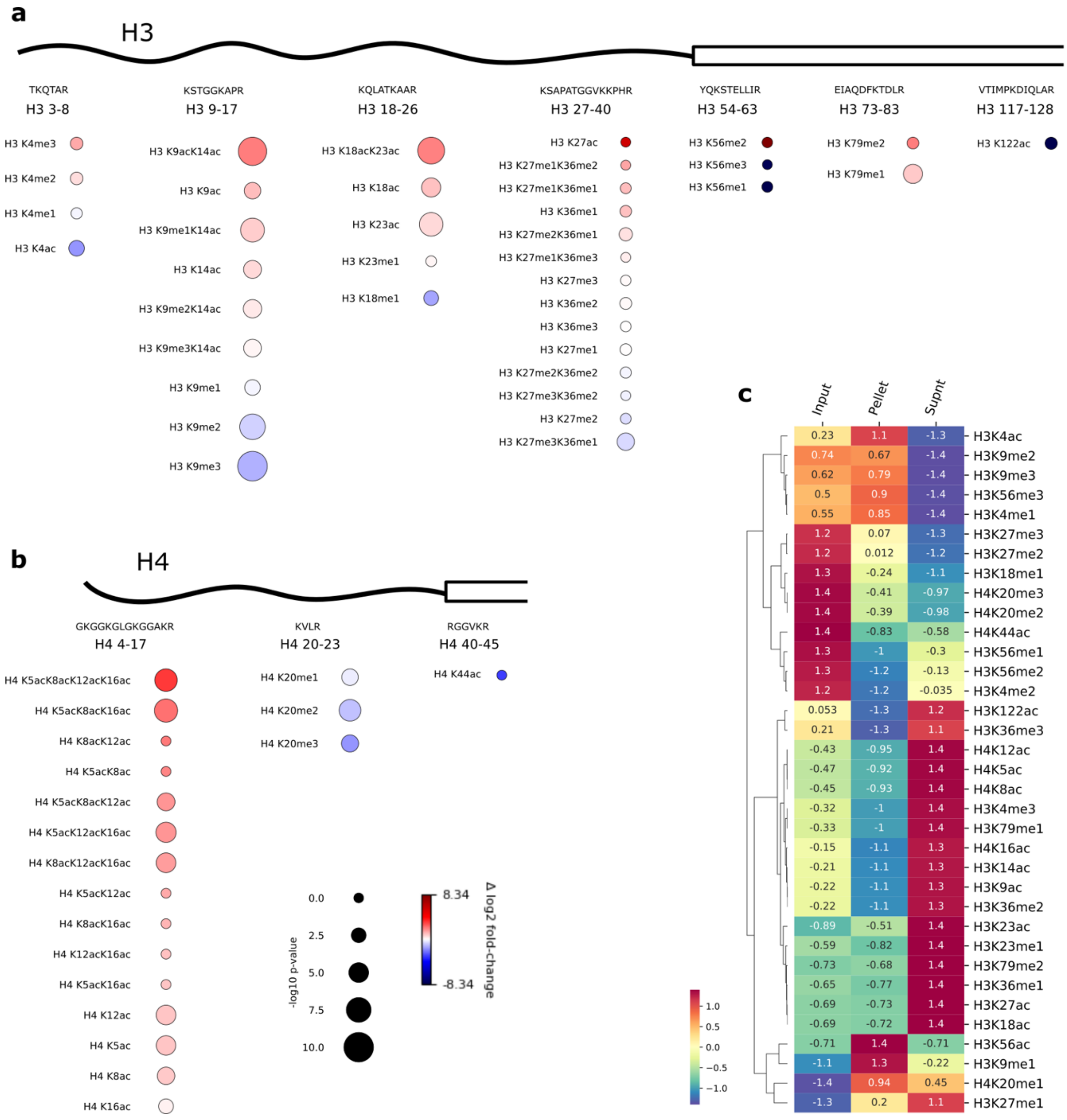


**Extended Data Fig. 7: Mass spectrometry identification of histone PTM marks with biased enrichment during native mononucleosome condensation experiments.**

**a**,**b**, Histone PTM marks detected in each histone H3/H4 peptides are shown. Its relative enrichment difference compared with the unmodified peptide is represented by color (red: more enriched in supernatant, blue: more depleted in supernatant) and its signification is represented by the size of bubble (-log p-value). **c**, Combinatorial histone PTM enrichment data was aggregated into single PTM modifications, and the relative enrichment in each phase of condensation (input/pellet/supernatant) is shown in the z-score heat map.

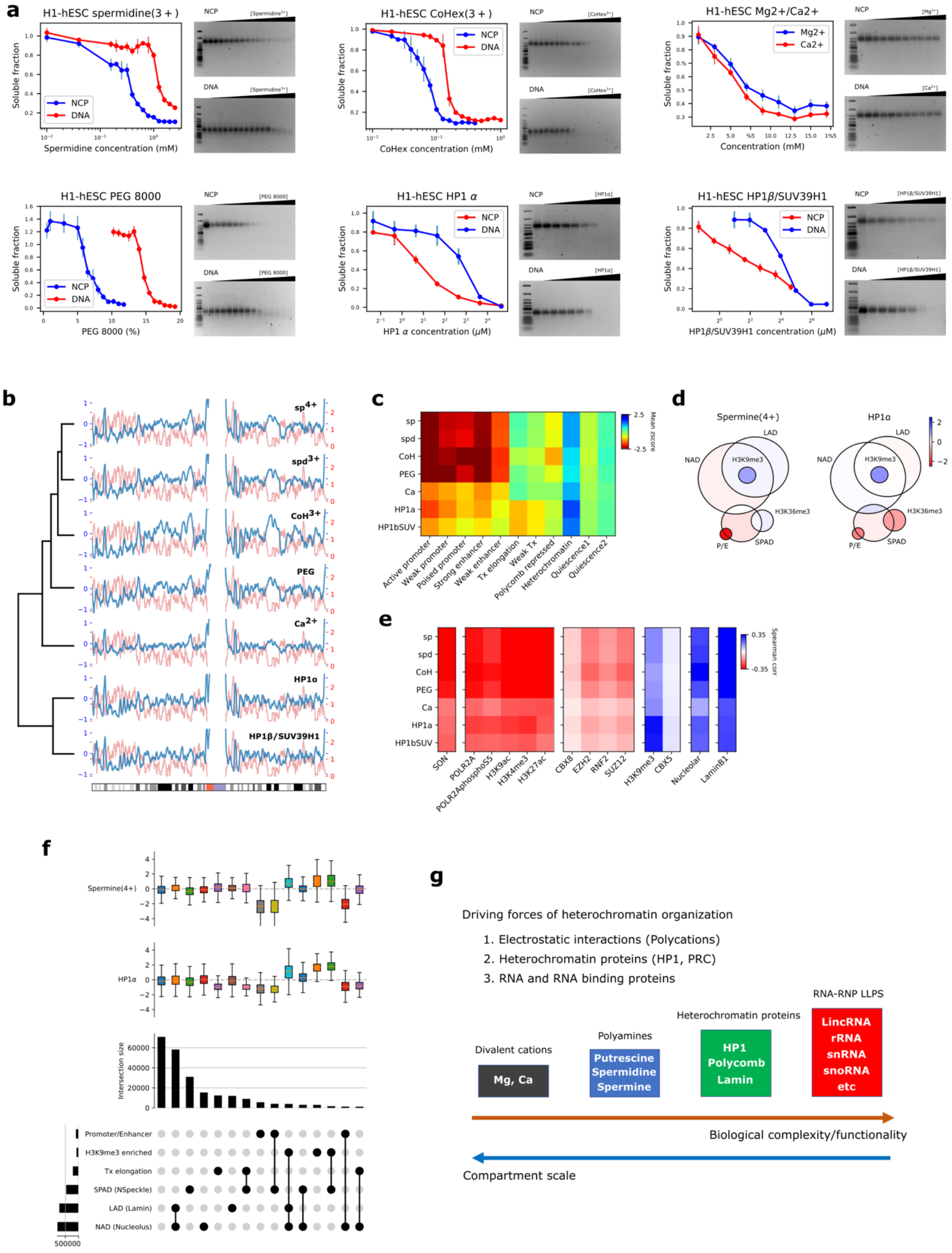

**Extended Data Fig. 8: Condense-seq of H1-hESC native mono-nucleosomes using various condensing agents.**

**a**, The soluble fraction of nucleosomes was measured by titrating the various condensing agents other than spermine, including spermidine, cobalt-hexamine, magnesium/calcium, PEG 8000, and HP1$\alpha$, HP1$\beta$ with SUV39H1 complex. **b**, Condensability scores were computed in 10kb resolution and plotted over chromosome 1 (blue) and compared with the gene expression level (red). Hierarchical clustering of the condensability profile shows that all ionic condensing agents (spermine/spermidine/cobalt-hexamine/PEG/calcium) are clustered together but other protein-based condensing agents (HP1$\alpha$ and HP1$\beta$) are clustered in a separate group. Comparison of condensability scores across various chromatin states (**c**) and various nuclear compartments (**d**, LAD: lamin-associated domain, NAD: nucleolar-associated domain, SPAD: nuclear speckle-associated domain, P/E: promoter or enhancer) for different condensing agents. **e**, Correlations between nucleosome condensability and the enrichment of other nuclear compartment hall mark proteins are represented as heat map. **f**, Upset plot of intersections between various nuclear compartments and the corresponding nucleosome condensabilities by spermine and HP1$\alpha$ shown as bar plots. **g**, Hypothetical hierarchal model of the biophysical driving force of chromatin organization: At a large scale, chromatin is compartmentalized via ubiquitous charge–charge interactions, but other proteins and RNAs are involved to generate sub-compartments that are smaller in scale but more specific function directed.

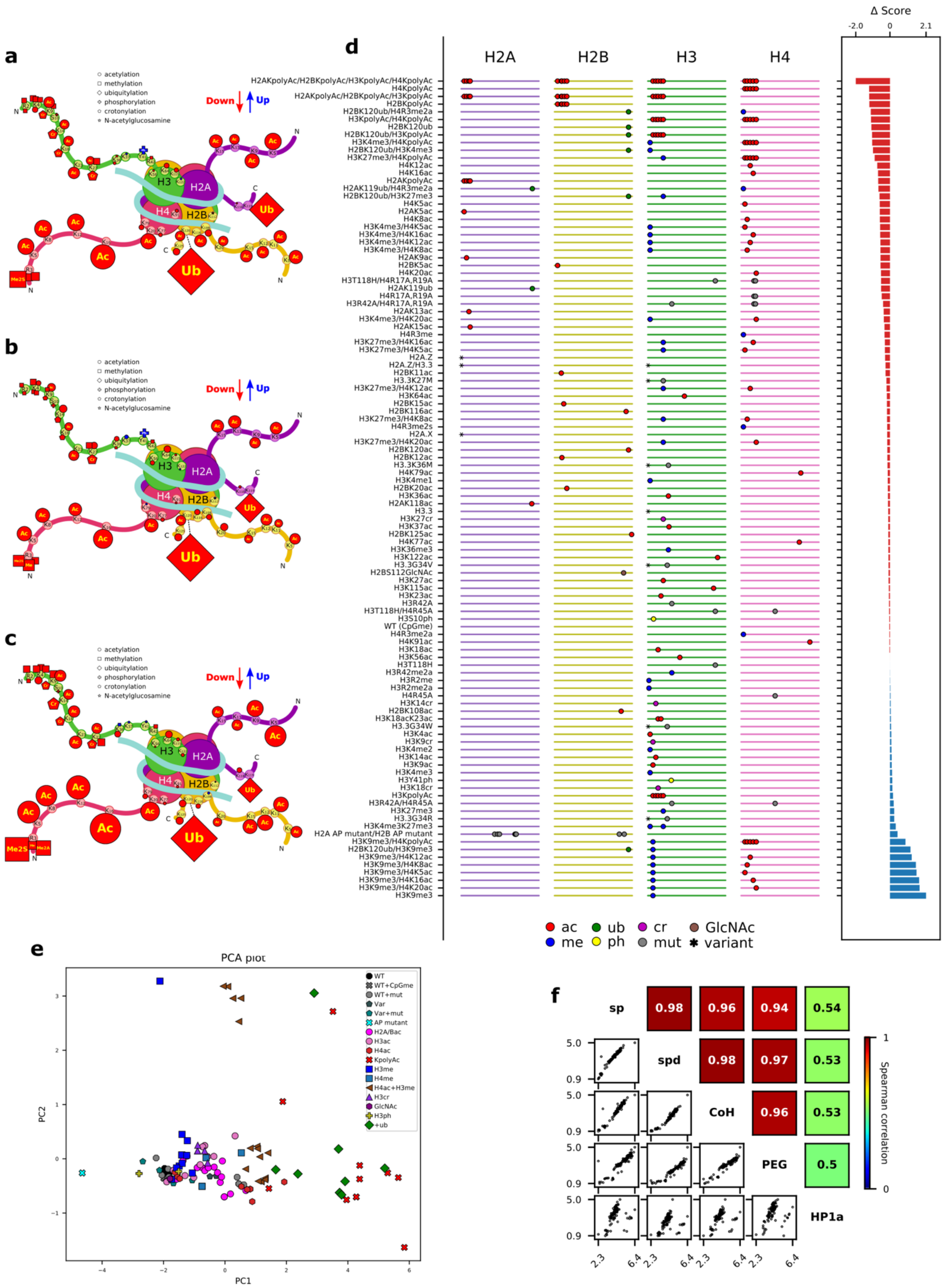

a
acetylation
methylation
ubiquitylation
phosphorylation
crotonylation
N-acetylglucosamine
Down
Up
H3
H2A
H4
H2B
Ac
Ub
b
c
d
H2A
H2B
H3
H4
Δ Score
-2.0
0
2.1
H2AKpolyAc/H2BKpolyAc/H3KpolyAc/H4KpolyAc
H4KpolyAc
H2AKpolyAc/H2BKpolyAc/H3KpolyAc
H2BKpolyAc
H2BK120ub/H4R3me2a
H3KpolyAc/H4KpolyAc
H2BK120ub
H2BK120ub/H3KpolyAc
H3K4me3/H4KpolyAc
H2BK120ub/H3K4me3
H3K27me3/H4KpolyAc
H4K12ac
H4K16ac
H2AKpolyAc
H2AK119ub/H4R3me2a
H2BK120ub/H3K27me3
H4K5ac
H2AK5ac
H4K8ac
H3K4me3/H4K5ac
H3K4me3/H4K16ac
H3K4me3/H4K12ac
H3K4me3/H4K8ac
H2AK9ac
H2BK5ac
H4K20ac
H3T118H/H4R17A,R19A
H2AK119ub
H4R17A,R19A
H3R42A/H4R17A,R19A
H2AK13ac
H3K4me3/H4K20ac
H2AK15ac
H4R3me
H3K27me3/H4K16ac
H3K27me3/H4K5ac
H2A.Z
H2A.Z/H3.3
H2BK11ac
H3.3K27M
H3K27me3/H4K12ac
H3K64ac
H2BK15ac
H2BK116ac
H3K27me3/H4K8ac
H4R3me2s
H2A.X
H3K27me3/H4K20ac
H2BK120ac
H2BK12ac
H3.3K36M
H4K79ac
H3K4me1
H2BK20ac
H3K36ac
H2AK118ac
H3.3
H3K27cr
H3K37ac
H2BK125ac
H4K77ac
H3K36me3
H3K122ac
H3.3G34V
H2BS112GlcNAc
H3K27ac
H3K115ac
H3K23ac
H3R42A
H3T118H/H4R45A
H3S10ph
WT (CpGme)
H4R3me2a
H4K91ac
H3K18ac
H3K56ac
H3T118H
H3R42me2a
H3R2me
H3R2me2a
H4R45A
H3K14cr
H2BK108ac
H3K18acK23ac
H3.3G34W
H3K4ac
H3K9cr
H3K4me2
H3K14ac
H3K9ac
H3K4me3
H3Y41ph
H3K18cr
H3KpolyAc
H3R42A/H4R45A
H3K27me3
H3.3G34R
H3K4me3K27me3
H2A AP mutant/H2B AP mutant
H3K9me3/H4KpolyAc
H2BK120ub/H3K9me3
H3K9me3/H4K12ac
H3K9me3/H4K8ac
H3K9me3/H4K5ac
H3K9me3/H4K16ac
H3K9me3/H4K20ac
H3K9me3
ac ub cr GlcNAc
me ph mut variant
e
PCA plot
WT
WT+CpGme
WT+mut
Var
Var+mut
AP mutant
H2A/Bac
H3ac
H4ac
KpolyAc
H3me
H4me
H4ac+H3me
H3cr
GlcNAc
H3ph
+ub
PC1
PC2
f
sp
spd
CoH
PEG
HP1a
0.98
0.96
0.94
0.54
0.98
0.97
0.53
0.96
0.53
0.5
5.0
0.9
2.3
6.4
Spearman correlation
1
0

**Extended Data Fig. 9: Summary of histone PTM effects on the nucleosome condensation by various condensing agents on the synthetic nucleosome library with PTM marks.**

The effects of single PTMs on nucleosome condensation are depicted by the cartoons (**a**: spermidine, **b**: cobalt-hexamine, **c**: PEG 8000 as the condensing agent). Each symbol represents different types of PTMs as shown in the legend, and the size is proportional to the strength of effects. The colors of the marks indicate the direction of the effect (red: decreases condensation, blue: increases condensation) compared with the unmodified control. **d**, All condensability scores of the PTM library using HP1α as a condensing agent are summarized in the ladder bar plot. The library members are sorted from the lowest to the highest condensability scores from top to bottom. On the left panel, the ladder-like lines represent each histone subunit peptide from N-terminal (left) to the C-terminal (right). Each mark on the line indicates the location of the PTMs and the shape of the marks represents the PTM type (ac: acetylation, me: methylation, cr: crotonylation, ub: ubiquitylation, ph: phosphorylation, GlcNAc: GlcNAcylation, mut: amino acid mutation, var: histone variant). On the right panel, differences in the condensability score compared with the unmodified control are shown as bar plots for each member of the library. **e**, PCA analysis was conducted by combining the condensability scores of all five condensing agents (spermine/spermidine/cobalt-hexamine/PEG 8000/HP1α) into the five-dimensional state vector. In the PCA plot, each member of the library is represented by a symbols according to categories such as canonical wild-type nucleosome (WT), wild type with CpG methylation (WT+CpGme), mutations on wild type (WT+mut), nucleosome with histone variants (Var), mutations on histone variants (Var+mut), Acidic patch mutants (AP mutants), and nucleosomes with acetylation on H2A/B dimer (H2A/Bac), acetylation on H3 (H3ac), acetylation on (H4ac), having poly-acetylation (KpolyAC), methylation on H3 (H3me), methylation on H4 (H4me), acetylation on H4 and methylation on H3 (H4ac + H3me), crotonylation on H3 (H3cr), GlcNAcylation (GlcNAc), phosphorylation on H3 (H3ph), and ubiquitylation (+ub), all of which are shown in the figure legend. **f**, Comparison of condensability scores across different condensing agents. Scatter plots of condensability across different condensing agents are shown in the lower triangle, and the corresponding Spearman's correlations are shown in the upper triangle of the matrix.

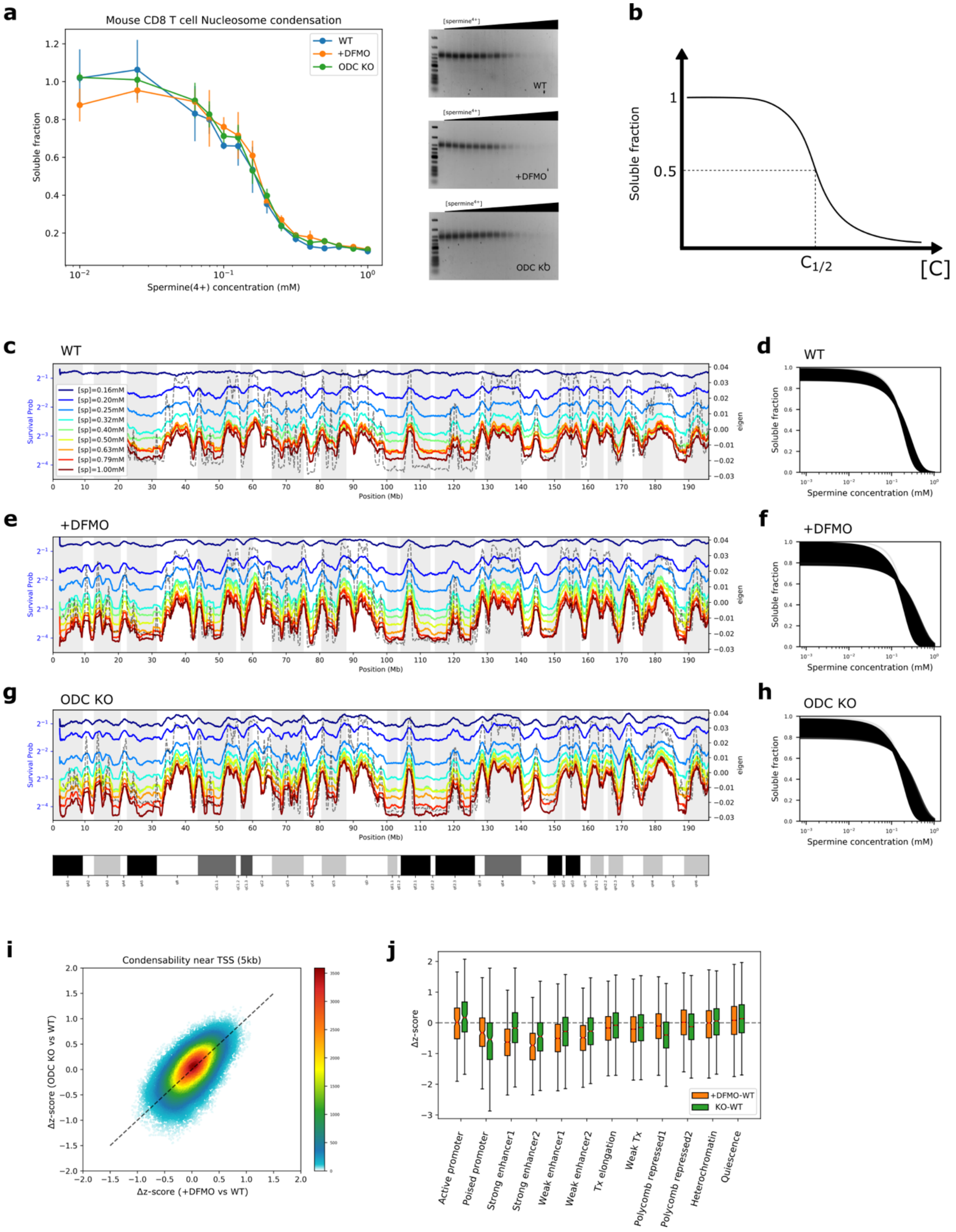


**Extended Data Fig. 10: Condense-seq measurements of nucleosomes purified from mouse CD8⁺ T cells.**

**a**, Soluble fractions were measured via UV-VIS spectroscopy and run in 2% agarose gel after condensation in various spermine concentrations. **b**, Condensation point ($c_{1/2}$) is defined by the concentration of condensing agent when the soluble fraction is half the input, so it is reversely correlated with condensability score. **c–h**, Soluble fractions of nucleosomes in various spermine concentrations were calculated and plotted over chromosome 1 in 10kb resolution. $C_{1/2}$ was computed for each bin after fitting the soluble fraction change with a logistic function as shown fitting curves of all bins (**d**, **f**, **h**), and polyamine deficient conditions show broader distribution of condensation points. (**c**, **d**: wild type control, **e**, **f**: +DFMO, **g**, **h**: ODC KO) **i**, The scatter plot of Δ z-score of condensability near TSS shows a high correlation between +DFMO and ODC KO. **j**, The Δ z-score of condensabilities is computed as the difference between the standardized condensability of +DFMO or ODC KO conditions and the wild type control and then categorized into the corresponding ChromHMM chromatin states.
